# Presence of a resident species aids invader evolution

**DOI:** 10.1101/2020.05.19.103978

**Authors:** Josianne Lachapelle, Elvire Bestion, Eleanor E Jackson, C-Elisa Schaum

## Abstract

Phytoplankton populations are intrinsically large and genetically variable, and interactions between species in these populations shape their physiological and evolutionary responses. Yet, evolutionary responses of microbial organisms in novel environments are investigated almost exclusively through the lens of species colonising new environments on their own, and invasion studies are often of short duration. Although exceptions exist, neither type of study usually measures ecologically relevant traits beyond growth rates. Here, we experimentally evolved populations of fresh- and seawater phytoplankton as monocultures (the green algae *Chlamydomonas moewusii* and *Ostreococcus tauri*, each colonising a novel, unoccupied salinity) and co-cultures (invading a novel salinity occupied by a resident species) for 200 generations. Colonisers and invaders differed in extinction risks, phenotypes (e.g. size, primary production rates) and strength of local adaptation: invaders had systematically lower extinction rates and broader salinity and temperature preferences than colonisers – regardless of the environment that the invader originated from. We emphasise that the presence of a locally adapted species has the potential to alter the invading species’ eco-evolutionary trajectories in a replicable way across environments of differing quality, and that the evolution of small cell size and high ROS tolerance may explain high invader fitness. To predict phytoplankton responses in a changing world, such interspecific relationships need to be accounted for.

## Introduction

Ecology and evolution affect invader success and native species’ responses, and in a warming, changing world, invasion scenarios are likely to become more frequent [1-3]. Eco-evolutionary studies on biological invasions for microbial populations are arduous to carry out - especially when encompassing an element of tracking evolutionary responses in real time for species less cultivable than bacteria (e.g. [4]), and are accordingly rare (but see e.g. [5,6])

When the survival of an organism – including invaders, colonisers, and resident species - hinges largely on physiological short-term responses, they can cope with a changed or changing environment through phenotypic plasticity. There, a given genotype produces a different phenotype in response to changes in the abiotic or biotic environment [7]. In asexually dividing microbes, these plastic responses occur within a single or a few generations and unless the new environment is lethal to most individuals in a sufficiently diverse population, it is unlikely that selection acts through sorting on such short time-scales. In clonal populations – the standard scenario in many laboratory experiments - evolution through *de novo* mutations is similarly rare on such short time-scales [8,9].

In the long term, the speed at which genetic sorting can affect the mean population trait in a diverse population depends on the amount of standing genetic variation, the strength of selection, and the size of the population [10,11]. Both plasticity and changes in allele frequency contribute to the magnitude and direction of an organism’s response to a novel environment. In aquatic microbes, short- and long-term responses to environmental change have been extensively studied in single isolate experiments (e.g. e.g. [12-15]). Far fewer studies consider ecologically complex environments [16-19]. Studies that have tested how species interactions evolve are traditionally carried out in environments that are of low quality for the focal species (traditionally low-nutrient or toxic environments), i.e. in environments that lead to slower growth and a drastic and near-fatal decline in population size of the focal species [8,20].

Under circumstances where resources are scarce, competition is often the main type of interaction [21-23]. Whether competition can then help or hinder evolution is highly context-dependent: Competition can explain changes in lineage or species frequencies as a function of the environment (e.g. light, nutrients, temperature) [24], but interactions between microbes need not be solely competitive and can range from competition to facilitation to mutualism to interdependence and combinations thereof [25-27], especially when environmental change is not leading to a decrease, but an increase in growth rate [28]. While the breadth of interactions is well understood in ecology [29-31], today’s models, when considering how microbes behave in a changing world, largely assume that interactions be competitive in nature [32]. Finally, these models are increasingly incorporating traits (and changes therein) other than growth rates, such as cell size and carbon fixation. Those traits are not routinely monitored in most selection experiments, where the focus tends to be on growth rates.

Here, we used an experimental evolution approach to conceptually investigate if coloniser or invader status changes an organism’s chances of survival in a new salinity and whether these dynamics differ between short-term (a few generations) and long-term (∼200 generations) responses under otherwise non-limiting conditions. We define colonisers as a species moving into a novel environment that is unoccupied (an ecologically rather unlikely scenario, but standard in most selection experiments), and invaders as species moving into a novel environment where local species are present. We used two green algae, the freshwater phytoplankton *Chlamydomonas moewusii* and the marine picoplankton *Ostreococcus tauri*. Both genera of green algae are cosmopolitan in natural systems and established model organisms for long-term studies in experimental evolution [33-35]. We evolved all populations for *ca* 200 generations either alone or in co-culture, in freshwater and marine conditions (Figure 1). To test whether differences between colonisers and invaders are general and repeatable in environments favouring faster growth, we superimposed a full-factorial temperature treatment, where all invaders and all colonisers were grown at 22°C (the culture ‘control’ temperature), 26°C (mild warming, and constituting a ‘better’, favourable environment where growth is faster and stress is lower), 32°C (extreme warming, constituting a ‘worse’, unfavourable environment, where growth is reduced and stress is higher), and a fluctuating temperature treatment, where temperature cycles between 22°C and 32°C every 3-5 generations (constituting a ‘better’, ‘favourable’ environment). The temperatures were chosen based on pilot studies examining the temperature tolerance curves of each species (see supporting information). We show that across all environments, freshwater and marine invaders fare better than colonisers, that samples selected under the invasion scenario evolve more generalist tendencies than samples selected as colonisers in the same environment, and that organisms in the invasion scenario evolve significantly different phenotypes compared to those in the coloniser scenario.

**Figure 1:**
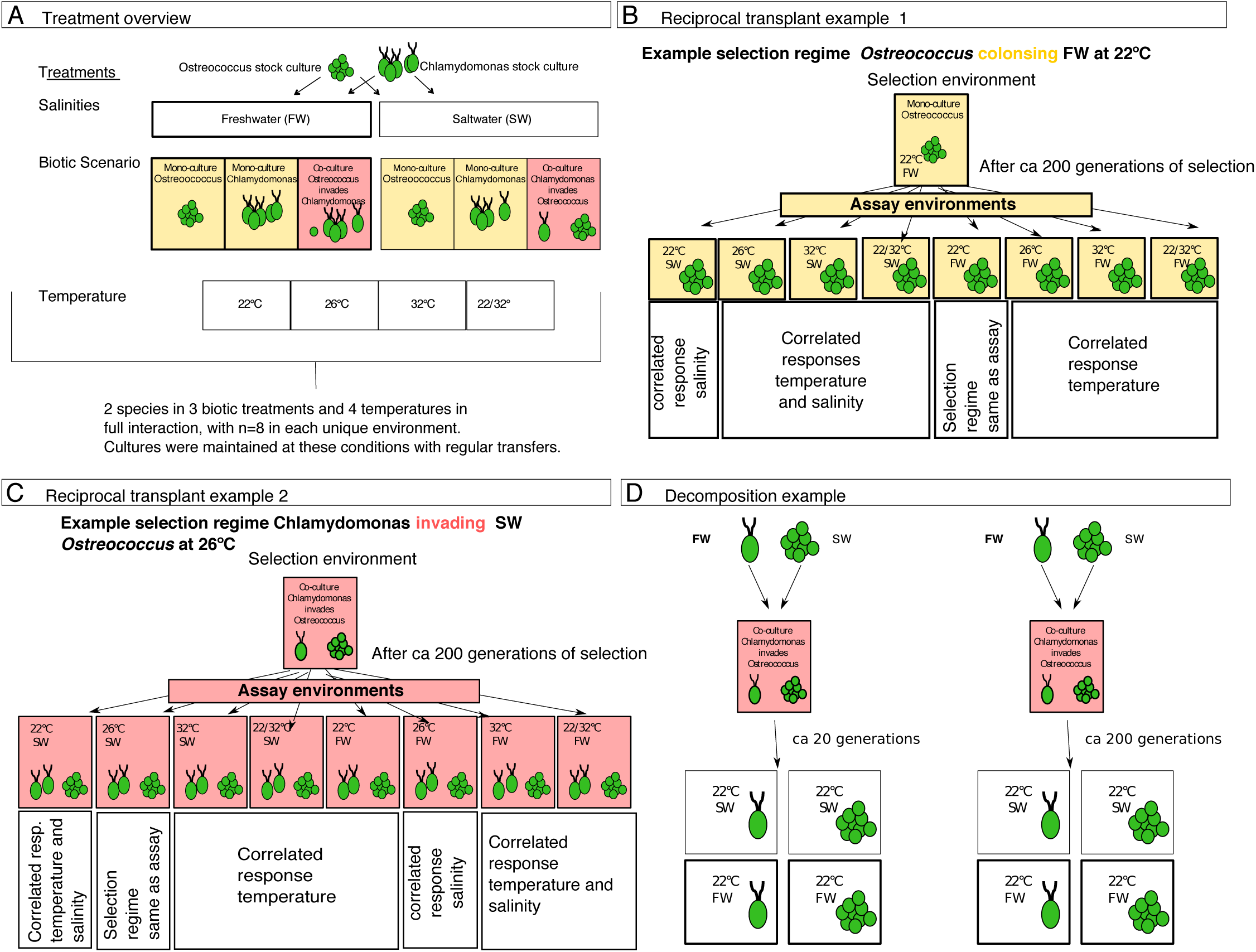
Experimental set-up. Throughout colour indicates the biotic scenario (yellow for colonisation, red for invasion), and the thickness of frames, the salinity (thick for freshwater and thin for salt water) **A) Treatment overview** Stock cultures of the marine *Ostreococcus* and the freshwater *Chlamydomonas* were used to inoculate two salinity regimes (a freshwater, FW, and a saltwater treatment, SW), crossed with three biotic experiments (*Ostreococcus* in monoculture, *Chlamydomonas* in monoculture, or co-culture with *Ostreococcus* invading *Chlamydomonas* and *vice versa*), and four temperature regimes (22°C as the control, 26°C as moderate warming, 32°C as extreme warming, and a variable environment where temperature cycled between 22°C and 32°C twice weekly, for a total of 24 unique selection environments. n=8 for each unique combination of salinity, biotic, and temperature regime. Samples were propagated weekly by batch transfer for approximately 200 generations. Light yellow denotes *Ostreococcus* or *Chlamydomonas* in mono-culture, i.e. the focal species is colonising. Light red denotes samples grown in co-culture. There, the focal species is invading. **B and C) example reciprocal assay** At the end of the experiment, all samples were assayed in all salinity and temperature treatments. **D) Decomposition of evolved samples:** Further, samples evolved in co-culture were decomposed into two separate monocultures and then assayed at both salinities at their selection temperature to test whether samples had evolved to depend on each other.

## Results

### Extinction risk is lower for invaders than for colonisers

Rapid changes in salinity can reduce the probability of phytoplankton survival and may limit population sizes to a point at which evolutionary adaptation becomes increasingly unlikely [36,37]. In our case, populations of all surviving cultures remained large enough to supply mutations and avoid drift at *ca* 10^5^ cells mL-1, which ensures a large enough supply of mutations for an evolutionary response, and makes differences between colonisers’ and invaders’ or different species’ responses in the long-term likely to be independent of population size.

Extinction rates in a new salinity were lower in invading than colonising phytoplankton (in a novel salinity, invading *Chlamydomonas* were two thirds less likely to go extinct than colonising *Chlamydomonas*, and for invading *Ostreococcus*, the likelihood of extinction was halved compared to colonising *Ostreococcus*, survival analysis: z = -2.90, P = 0.0037 Supporting Tables 1 and 2). Extinction events occurred early on in the experiments (within the first 70 generations, z = -4.13, P = 3.6 × 10^−5^; Fig. 2; Supporting Table 1 and 2, Supporting Figure 1), with no further extinctions after 100 generations. After the first ∼ 70 generations, population sizes in all surviving invader cultures stabilised and were no longer statistically different from colonisers (F _1,2_ =1.72, p = 0.32; more details in Supporting Tables 3 to 4).

**Figure 2:**
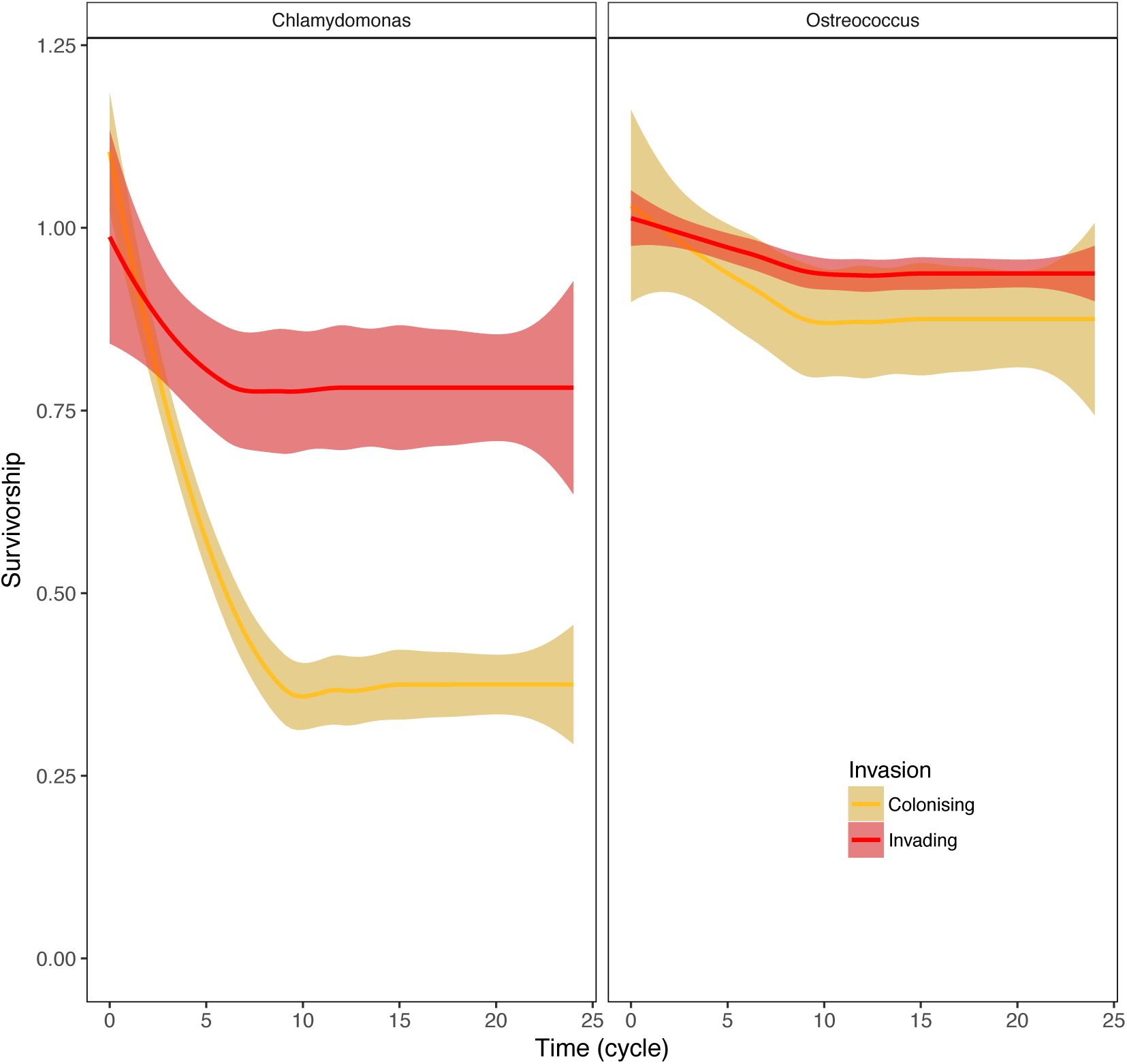
Survival when invading into a novel salinity from rare is enhanced in the presence of a resident species across all selection regimes. Displayed is the mean survivorship for samples in all temperature regimes over time (where one cycle or transfer corresponds to one week), with 1.0 as 100% o populations surviving, and shaded areas denoting 95% confidence intervals. All treatment combinations started with N = 8. The proportion of populations and replicates surviving decreased rapidly at the beginning of the experiment but levelled off after about 10 transfers as extinctions stopped. The number of surviving biological replicates at the end of the experiment is shown in Table S1. Invader trajectories in red, coloniser trajectories in yellow.

70 -100 generations is a time-frame corresponding roughly to a single growing season of fast-growing green algae, and is comparable to other marine microbial experiments where evolution occurred on the scale of just below 100 to a few 100 generations [13,15,38], though as few as two generations have been reported to suffice for an evolutionary response [39].

Theory predicts that evolutionary potential should be high in good quality environments leading to an increased or unchanged fitness, and that extinction risk should be low in environments that fluctuate predictably (e.g. [40]). We found that extinctions were indeed overall lowest in the ameliorated environments, i.e. under mild warming at 26°C, and in the fluctuating environment (survival analysis: z = -1.22, P = 0.043; Supporting Figure 1, Supporting Table 2). *Ostreococcus* colonisers had a small but significant (see Supporting Table 2) advantage over *Chlamydomonas* colonisers, with higher survivability over all. While we cannot determine the mechanism by which co-culture favours survival, or whether *Ostreococcus* are per se better colonisers than *Chlamydomonas* (Fig. 2 and Supporting Table 2), we can begin to quantify the effects of abiotic environment and species interactions on evolutionary and short-term responses.

### Invaders and colonisers differ in salinity and temperature tolerances

Survival is insufficient: Once extinction is no longer one of the main mechanisms driving responses, we need to know the performance of the population across environments, and the phenotypes they might display. To integrate plastic and evolutionary responses into ecosystem [41] and individual-based models [42], and to better understand the dynamics in laboratory experiments [43] knowledge of phenotypic traits and organismal biology is needed. Here, colonisers differed from invaders in the magnitude of their evolutionary response, their ability to grow in their ancestral environments, and in the phenotypes they evolved.

When colonisers were transferred back into their ancestral salinity, their growth rates were the same as or lower than they had been in that same salinity before evolution in a novel-salinity environment (e.g. average growth rate per day of the coloniser in the novel salinity after evolution in novel salinity: 1.13 ± 0.02 SEM, and after transfer back into the ancestral salinity: 0.79 ± 0.02 SEM, Fig. 3, Supporting Tables 5 and 6 for more details). Growth rates of invaders were overall higher, and did not decrease significantly upon being transplanted back into their ancestral salinity (average growth rate of the invader in the novel salinity after evolution in novel salinity: 1.31 ± 0.01 SEM, and after transfer back into the original salinity:1.29 ± 0.01 SEM). Invaders and colonisers also differed with regards to their responses to warming (supporting Tables 7 and 8 for details), where invaders again outperformed colonisers. This pattern was exacerbated under mild warming in 26°C where invader growth rates were on average 1.3 times higher than coloniser growth rates (tukey post hoc, P<0.001), and under the fluctuating treatment, with an average fold increase of invaders *vs* colonisers of 1.2 (tukey post hoc, P<0.001). The most pronounced advantage of invaders over colonisers was at the selection temperature (Fig. 4 A, Supporting Tables 7 and 8). Growth rates were diminished under environmental deterioration at 32° C, and this decrease in growth was the least pronounced in invading species (Fig. 4A). We find support that in the unfavourable environments (high temperature, changed salinity), intracellular reactive oxygen species (ROS) production is higher, and ROS tolerance impeded, but that this effect is more pronounced in colonisers than invaders (Supporting Figures 2 and 3).

**Figure 3:**
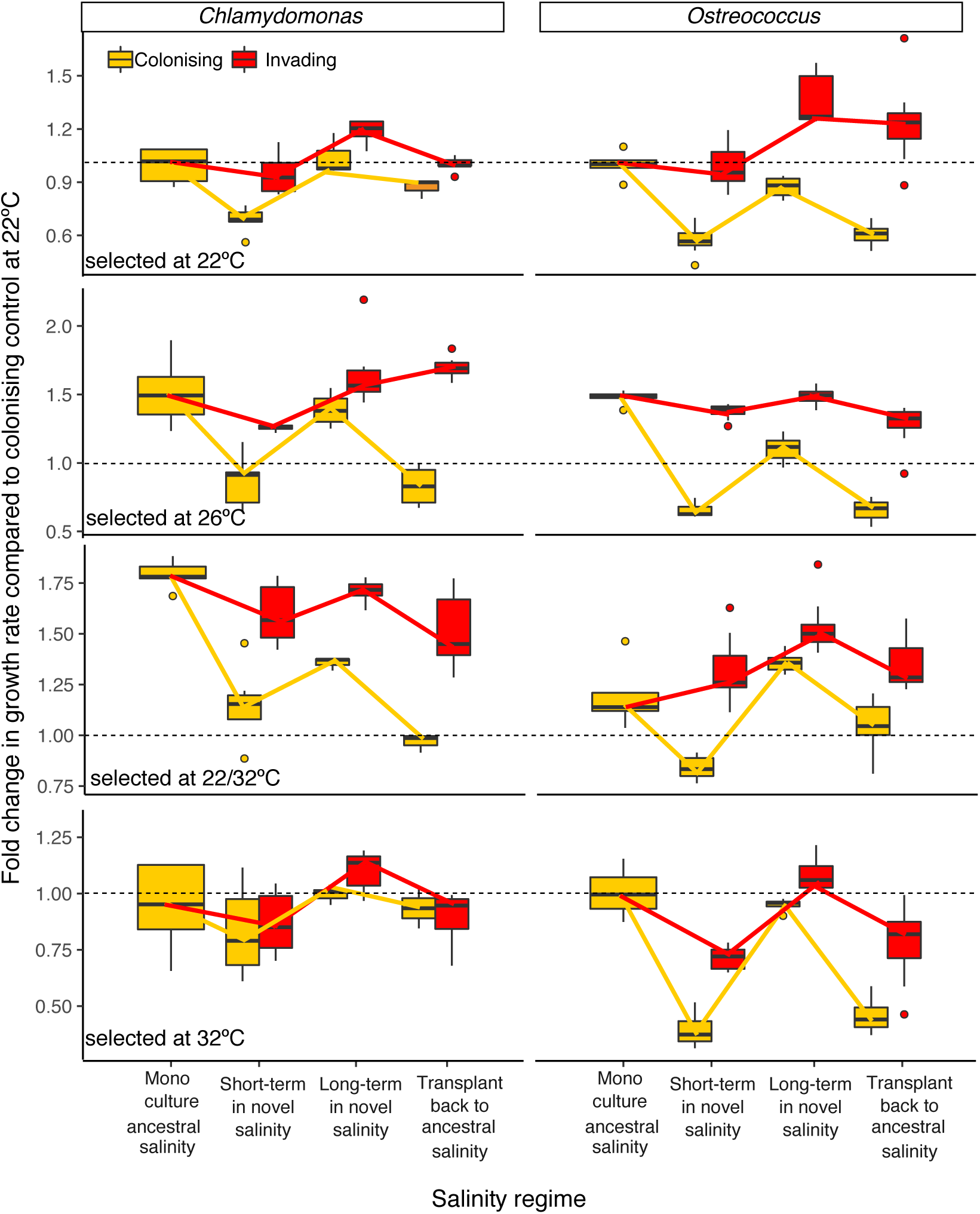
Invaders and colonisers differ in their responses to salinity regimes across time scales. Salinity regime ‘Mono-culture ancestral’ denotes that the species was assayed in its ancestral salinity in mono-culture after evolution in the ancestral salinity in mono-culture. ‘Short-term in novel salinity’ is for growth rates measured after the sample had spent two transfers in the novel salinity. ‘Long-term in novel salinity’ is the growth rate of the sample in the salinity that it was evolving in. ‘Transplant back to ancestral is for growth rates measured when samples were transferred back into the ancestral environment after evolution in the novel salinity. We express changes in growth as compared to *Ostreococcus* or *Chlamydomonas* in its home salinity at 22°C in mono-culture, i.e. values < 1 (below dotted line) indicate that a sample grew more slowly than the same species in mono-culture, in its home salinity at 22°C, and values > 1 (below dotted line) indicate that they grew faster. Each panel is for one selection temperature. Orange boxplots are for colonising species, and red, for invaders. See Table S2 for details on n per treatment. Boxplots are displayed as is standard, with the belt indicating the median. Fitted lines are for visualisation.

**Figure 4:**
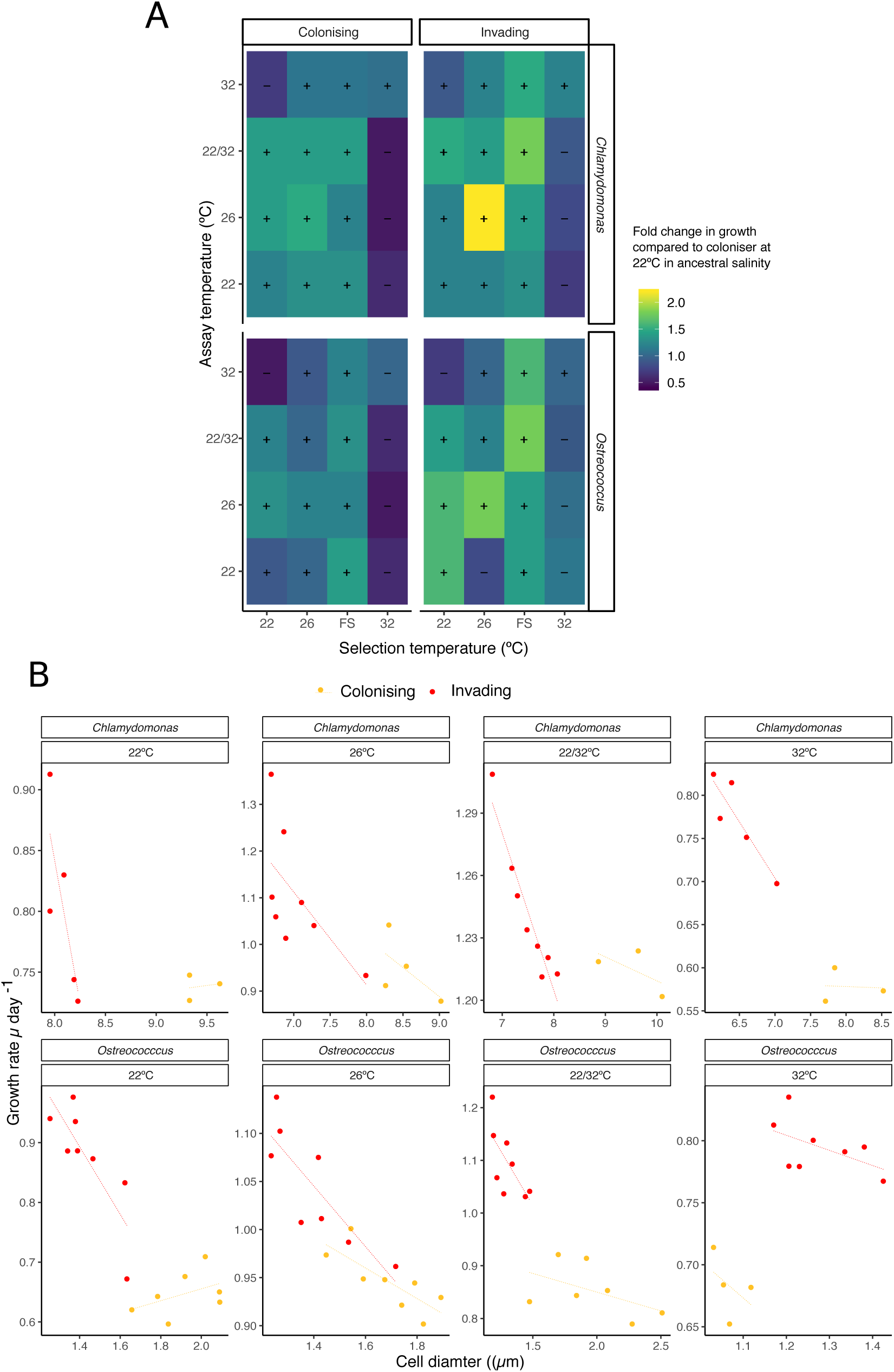
Invaders fare better than colonisers in deteriorated (e.g. 32°C degree) and ameliorated (e.g. 26°C) environments. Invaders evolve small cells, yielding higher growth. Visualisation of reciprocal temperature assays for colonisers and invaders. A: Tiles indicate whether the response to the assay condition (y axis) of invading and colonising cultures selected at a specific temperature (x axis) was to grow more slowly (purple hues) or faster (green and yellow hues) than the coloniser at 22°C in the ancestral salinity. B: Within each regime (invasion –red, or colonisation – orange), invaders tend to have smaller cells. Smaller cell size is associated with faster growth (and higher ROS tolerance, as well as lower ROS production, see Supporting Figure 11). Number of replicates varies due to treatment specific differences in extinction probability.

### Phenotypic traits of colonisers and invaders

Cell size overall declined with selection temperature regardless of selection regime or species. *Ostreococcus* was more reactive to temperature than *Chlamydomonas* overall (Supporting Figure 4, Supporting Tables 9 and 10), and whether the species was invading or colonising also had an impact on the focal species’ cell size (Supporting Tables 9 and 10, Supporting Figure 4), with smaller invaders than colonisers. Cell size of *Chlamydomonas* was more likely to change in response to a resident species than cell size of *Ostreococcus*, with *Chlamydomonas* cells up to 1.43 fold smaller after evolution invading the marine species in saltwater than after colonising saltwater on their own (also Figure 4B).

While we cannot disentangle the relative contributions of the individual species to Net Photosynthesis rates in the co-cultures (NP, i.e. rates of photosynthesis after respiration has been accounted for), net-photosynthesis per gram carbon of evolved co-cultured samples was on average 13% higher than expected from the NP of the same two species at the same salinity in monoculture in line with over-yielding observed in other species [44] (Supporting Figure 5, and Supporting Tables 11 -14). The same pattern emerged when we assayed the same species at the same salinity after decomposition of the co-cultures into monocultures (e.g. physically separating a former mixed culture of *Chlamydomonas* residents and *Ostreococcus* invaders into monocultures (Supporting Tables 11 -14)).

### Experimental community decomposition

Experimentally separating (‘decomposing’) the evolved co-culture samples into monocultures yielded insights into how strongly the invaders had adapted to the presence of the resident species, and what effect the invader had on the growth of the resident species (see methods for details). In samples that had only lived in co-culture for two transfers (<20 generations), growth after decomposition was indistinguishable from growth in mono-cultures at the same salinity and temperature (Supporting Table 15, Supporting Figure 6). In contrast, in samples that had lived in co-culture for ∼200 generations, growth of the invading species when assayed alone in the selection salinity was reduced by up to 30% compared to when assayed in co-culture (Supporting Figure 7), and compared to the same species evolved in mono-culture at the same salinity/temperature regime. Of the resident species, *Ostreococcus* selected in co-culture with *Chlamydomonas* showed evidence of a marked decrease in growth when the invading *Chlamydomonas* was removed. Resident *Chlamydomonas* grew faster when the invading *Ostreococcus* population was removed, with no significant effect of temperature on this pattern.

In the decomposed samples, patterns in net primary production of former invaders mirrored the patterns found in growth rates: invaders always photosynthesised less after decomposition than the same species evolved in monoculture in the same selection salinity. High photosynthesis rates in formerly invaded *Chlamydomonas* were in line with higher growth rates in formerly decomposed *Chlamydomonas* (Supporting Tables 11-14, Supporting Figure 7). The resident species *Ostreococcus* photosynthesised more after decomposition than when evolved in monoculture - but grew more slowly. The higher NP rates were, at least for the duration of the assay (two weeks), not directly channelled into growth, indicating that the presence of other species may explain hitherto often observed but poorly explained variations in growth rates in more complex systems (but see [16,45]). We found that samples with the highest surplus NP (or least increase in growth) had a tendency to have higher Nile Red fluorescence, indicating higher lipid storage (Supporting Figure 8). Similar responses including high rates of NP but suppressed growth can be achieved by merely spiking *Ostreococcus* and *Chlamydomonas* cultures with water conditioned by the other species (Supporting Figure 9).

## Discussion and Conclusions

Rapid adaptation to a novel salinity or the evolution of salinity tolerance are major driving forces in determining the distribution and phenotypic characteristics of phytoplankton communities [46-50]. Changes in salinity (IPCC, 2014), particularly in combination with elevated temperatures, have the potential to impact the phenotypic characteristics of phytoplankton species, the communities they populate, and the role of phytoplankton species on aquatic food webs and global nutrient cycles [48,50,51]. Here, we found that species *colonising* a new salinity were prone to extinctions, but that survivors rapidly became locally adapted to their novel salinity. Rapid evolution to a novel salinity has been proven before [34,52], but evolution as a single species might not be a common ecological scenario, as species are likely to not arrive in a new environment and find it unoccupied. Species *invading* a new salinity were less likely to go extinct and evolved high tolerance to both fresh and saltwater, especially under environmental amelioration, such as mildly elevated or rapidly fluctuating temperatures. Invaders had higher survival and growth rates, and were also characterised by overall smaller cell size, lower reactive oxygen species (ROS) production, higher ROS tolerance, and a tendency to store lipids. ROS are a natural by-product of cellular metabolism, but can damage the cell at high quantities [53]. Therefore, higher tolerance toward or lower quantities of ROS may infer a fitness benefit [54]. Taking into account growth rates across all temperature treatments, the invader became more of a generalist, with better performance across multiple environments. Successful invading species often have traits associated with generalists (see e.g. [55,56]), but whether generalist traits enable successful invasions or whether organisms evolve to have more generalist traits as a consequence of invasions remains an open question.

Our results suggest that under warming and increased climate variability, invasions through small, warm-adapted taxa with intrinsically elevated metabolic and growth rates may become more frequent (‘tropicalisations’, see [57,58]), with nigh-unpredictable consequences on aquatic ecosystems as a whole. As changes in fitness, cell size and metabolic activity are often linked [59,60], it stands to reason that one possible mechanisms for higher invader fitness in our invader samples lies in their ability to rapidly down-regulate cell size [61-63], which in turn might be what is giving rise to their ability to better handle reactive oxygen species [54,62] (Figure 4B and Supporting Figure 10 – the smallest cells had highest fitness and were better able to detoxify ROS). The dynamics and mechanisms of increasing fitness under constant directional selection are well understood [64,65], and experimental evolution lends itself well to linking environmental cause to evolutionary effect, but it is limited in accurately deciphering the mechanisms that underlie trait evolution. Strategies that increase fitness can vary over time [9] and when there are multiple genotypes in a population, evolutionary trajectories, as well as the traits evolved will depend on the environment as well as the genotype [39]. Due to the complex nature of fluctuating selection regimes, barring further analyses, for example on the level of the transcriptome, we cannot with certainty elucidate the exact mechanism that allows for the evolution of these strikingly different phenotypes in invasion *vs*. colonisation scenarios. Still, *Ostreococcus* selected in co-culture with *Chlamydomonas* showed evidence of a marked decrease in growth when the invading *Chlamydomonas* was removed, suggesting that interactions were mutualistic or facilitating in nature. *Chlamydomonas*, when *Ostreococcus* were removed, did *not* show a marked decrease in growth, making it seem likely that the fact of being an invader had direct phenotypic and fitness consequences regardless of the nature of the interaction.

Understanding the impacts of environmental change over evolutionary timescales will require that we experimentally investigate the mechanisms underlying the differences between colonisers and invaders, the direct effects of rising temperatures on species interactions, and the indirect reciprocal feedbacks between ecological and evolutionary dynamics [29,30,66-68].

## Supporting information

All Supporting Information

## Acknowledgments

This study was funded by a British Ecological Society ‘small grant’ to ES (BES 5705 / 6749). *Ostreococcus* samples were kindly provided by Samuel Barton and Sarah Heath. *Chlamydomonas* samples were sourced by EB. Pilot studies were carried out by ES and JL at the University of Edinburgh, where Nick Colegrave and Sinéad Collins provided bench space and equipment. Gabriel Yvon-Durocher provided bench and incubator space for the main experiment at the Environment and Sustainability Institute in Penryn, UK. The authors thank Etienne Low-Decarie and Georgina Brennan for comments that greatly improved the manuscript, and Stefanie Schnell for maintaining the cultures in Hamburg.

## Author contributions

ES and JL conceived and designed the experiment and wrote the manuscript. ES, EB, and EJ carried out experiments, and ES supervised laboratory work. JL and ES analysed data, and all authors contributed to writing the manuscript.

The authors declare no conflict of financial or other interest, and all data will be made available on zenodo or data dryad upon acceptance. During the pre-print stage data are available from the authors upon request.

## Methods

**Should the methods exceed the allowed number of pages, we will provide a methods summary here, and the detailed methods, in the supporting information.**

### Algae strains

The marine picoplankton *Ostreococcus tauri* (clone of the original OTH95) and the freshwater alga *Chlamydomonas moewusii* (CCAP 11/5B) were sourced as non-axenic stock cultures from the Roscoff culture collection and the CCAP (Culture Collection of Algae and Protozoa) respectively. The fact that these two species do not usually co-occur in nature is not problematic here because species invasion is the result of a new species being introduced into a new environment with resident species it has not interacted with before. Pilot studies revealed that long-term growth was impoverished upon removal of the associated bacterial component or after antibiotic treatment, and thus no further attempts at using axenic cultures were made for the purpose of this study. The total amount of bacterial co-inhabitants was tracked and did not change throughout the experiment. Samples were maintained in semi-continuous batch culture (i.e. a fixed volume of exponentially growing cells was transferred into fresh medium at regular intervals) at 22°C, 100µmol quanta*s^-1^*m^-2^ under a 12:12 hour light:dark cycle in INFORS ™ multitron incubators with integrated shakers until use. *Ostreococcus* was grown at a salinity of 32 (PSU, roughly 30g NaCl*l^-1^; referred to from now on as saltwater or SW) in f/2 media [69], *Chlamydomonas* in modified Bold’s media (roughly 0.025g NaCl*l^-1^; referred to from now on as freshwater or FW). Concentrations of major nitrogen and phosphorus sources were the same in the fresh- and saltwater media.

### Selection experiment

We set up our experiment using two salinity regimes (saltwater and freshwater, where we refer to the salinity that the species originated from as the ‘ancestral’ salinity) and three biotic regimes (two monoculture, and one co-culture scenario; Figure 1A). Residents are species evolved in their ancestral salinity (i.e. *Chlamydomonas* in freshwater, *Ostreococcus* in saltwater) either in mono-culture or in co-culture with an invading species. Invaders are species evolved in a novel salinity where the resident species is present (i.e. *Chlamydomonas* invading *Ostreococcus* in saltwater, *Ostreococcus* invading *Chlamydomonas* in freshwater). Finally, colonisers are species evolved in a novel salinity as a mono-colture (i.e. *Ostroeoccus* in freshwater, and *Chlamydomonas* in saltwater).

In pilot studies, we characterised temperature reaction curves for each species at each salinity. Temperature/salinity combinations that lead to a significant decrease in growth rate in the short-term compared to the coloniser in its ancestral salinity at 22°C were called ‘low’ quality or ‘unfavourable environments. Thermal environments where growth rate increased were called ‘high’ quality or ‘favourable’ environments (here, these are the 26°C and fluctuating environment – this is also reflected in samples’ abilities to deal with reactive oxygen species). Based on these pilot studies, the long-term experiment was replicated across four different temperature regimes, for a total of 24 unique treatments (x 8 biological replicates = 192 cultures, Figure 1A). The temperature regimes consisted of a fluctuating temperature treatment and three stable temperatures, encompassing a stable ambient 22°C treatment (control), a stable 32°C treatment (severe warming), and a stable 26°C treatment (mild warming). In the fluctuating temperature treatment, temperature was switched between 22°C and 32°C .ca every 3-5 generations.

We expect the first adaptive step to occur more rapidly in genetically diverse starting populations than in clonal populations, and have leveraged this in our study by starting with genetically diverse rather than clonal populations. [70]

All cultures started out as mono-cultures before invading species were added. Cultures were grown on 48-well plates with sterile, breathable membranes (Aeraseal ™, Sigma-Aldrich) to minimise uneven evaporation and air exchange across plates. Monocultures were initially inoculated with 100 cells of *Chlamydomonas* or 1000 cells of *Ostreococcus* to account for the difference in cell size. In co-cultures, the resident species were inoculated at 100 fold the biomass of the invading species, for an ‘invading from rare’ scenario at the beginning of the experiment. The invasion event occurred only once at the beginning of the experiment, after which we tracked the fate of the invaders throughout the experiment. The 48-well plates were positioned randomly in the incubator, and their position was changed every other day to minimise location effects. Cultures were maintained in semi-continuous batch culture, where well-mixed samples of 200µl were serially transferred into 1200µl of new media every 7-10 generations (‘transfers’). At each transfer, cell count was determined using an Accuri c6 (BD Scientific) flow cytometer at high flow rate. Cells from the two species grown in co-culture could be distinguished based on the SSC (side scatter for granularity), FSC (forward scatter for cell size), and FL3 (red fluorescence for chlorophyll content) channels (Supporting Figure 11), allowing for species growth curves to be tracked separately. To analyse differences in cell sizes between treatments and species, we calibrated the flow cytometer with beads of known size.

Cell counts at the beginning and end of each transfer cycle were used to calculate the rate of increase in cell numbers and approximate generation times. Rates of increase in cell number were determined assuming logistic growth (based on pilot experiments), using the formula

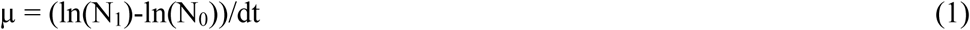

Where N_1_ is the cell count at the end, and N_0_ at the start of the transfer, and dt is the length of the transfer cycle (seven days).

The experiment was carried out for approximately 200 generations.

### Reciprocal assays

After 27 transfers in their respective selection environments, all samples were subjected to a full reciprocal transplant assay in all salinity and temperature regimes to test whether the surviving colonisers and invaders had adapted to the novel salinity in each temperature regime (Figure 1 B and C), and to calculate the magnitude of the short term and evolutionary responses (Figure 1B and C). A well-mixed sample from each surviving population was used to seed the assays. Assays were performed using the same inoculum size and duration of transfer cycle as during the selection experiment. The assays consisted of two transfers, where the first was used to allow the cultures to acclimate to the environment, and the second was used to measure the rate of increase in cell number as a proxy for fitness. Samples evolved at 22°C in their ancestral salinity in mono-culture were used as ‘evolved controls’, which take into account any evolution that may have occurred due to laboratory conditions *per se*.

We measured three types of responses: the short term response (occurring largely through rapid sorting and physiological acclimation within the same or a few generations, here, less than 10-14 generations), the long term response (likely largely evolutionary, > 100 generations), and the correlated response (growth in environments other than the selection environment, 10-14 generations after termination of the long-term experiment). See e.g. [38] for calculation of the magnitude of short-and long term responses, as well as responses in the reciprocal environments.

### Experimental decomposition of populations grown and evolved in co-culture into mono- cultures

To assess whether invaders evolved in co-culture had adapted to the novel salinity, the presence of the resident species, or both, we passed all co-cultured samples through a 5µm nitrocellulose filter, allowing *Ostreococcus* cells to pass, while *Chlamydomonas* cells remained on the filter, from which they could be rinsed off. Samples were then inspected under the microscope and flow cytometric data was again used as above to ensure a good separation of the two species. Samples were grown for two transfers in both their evolved and their ancestral salinity (Figure 1D). We compared the increase in cell number when they were grown on their own after decomposition to when they were grown in co-culture or had been selected for growth in monoculture. For logistic reasons, samples were only assayed at the temperatures that they had evolved in and not across all temperatures. To measure the short-term acclimation response to encountering another species, we re-created the starting conditions of the invasion experiment using mono-culture evolved samples (colonisers evolved in their ancestral salinity). These samples were inoculated to recreate the invasion from rare scenario as described above. The new co-cultures were maintained for two cycles, and then separated again by filtration afterwards. This tested for whether dependence on the species was established within very few generations (Supporting Figure 6).

### Characterisation of net primary production

To characterise phenotypic changes in the different treatments, we gathered data on cell size and chlorophyll through flow cytometry (Accuri B6), and measured rates of oxygen evolution and consumption using a 24-channel PreSens Sensor Dish Reader. For all phenotypic characterisations, samples were harvested during exponential growth. The reader was placed in the incubator at assay temperature in a manner such that the light gradient across the reader plate was minimal (<5 µE m^-2^ s^-1^). Glass vials were filled to 1.2 mL with the respective sample, i.e. colonisers, invaders, or decomposed samples, covered with para-film and sealed tight. The samples were then left in the dark for 35 minutes, and gently inverted before measurements of oxygen evolution at the light level in the incubator for 5 minutes, and measurements of oxygen consumption in the dark for another 5 minutes. A vial containing filtered Bold’s medium or f/2 medium at the appropriate salinity was used to account for any drift in the oxygen measurements. Cell count was determined using a flow cytometer as described above. The rates of oxygen evolution and consumption were then calculated per unit biomass, assuming spherical cells and carbon conversion factors after [71] (Supporting Figure 12 and Supporting Tables 16 and 17 for effects of selection regimes on biomass).

### Nile Red stain

A Nile Red stain was used as a proxy to determine relative quantities of intracellular polar and neutral lipids [72]. It works well for Ostreococcus [73] and while stains of the BODIPY class are preferred for quantification of lipids in Chlamydomonas, Nile Red can serve well to establish relative differences[74]. The dye was added to each 200 µL sample on a 9-well plate for a final concentration 15 m and left to incubate in the dark for 30 min, as pilot trials had shown that after this, fluorescence levels were stable long enough for the time taken to measure one 96 well plate. As Nile Red excites in the same wavelength as chlorophyll (FL3) and chlorophyll derivatives (FL2), samples were measured before and after adding the dye, and the chlorophyll fluorescence subtracted from the fluorescence obtained after staining the sample (Supporting Figure 9).

### ROS assay

We tested how capable samples were of detoxifying harmful reactive oxygen species (ROS) and also estimated the intra-cellular ROS levels in order to gain an estimate on whether samples under unfavourable conditions experience more stress, and are therefore producing more/ being less able to detoxify ROS. We used the protocols established by [54,62]. Samples from ‘unfavourable environments’ had higher intra-cellular ROS content, were less well able to detoxify harmful ROS (Supporting Figures 2,3, 11).

### Statistical analyses

All data were analysed in R versions 3.3.1 and 3.3.3 [75].

### Survival analysis

We first analysed the extinction dynamics by performing a survival analysis using a Cox proportional hazards regression model with the R package ‘survival’(Supporting Table 1 and 2). The model included biotic regime, temperature regime, and species as fixed effects. Biological replicate strains (per species) were treated as random effects. We also included a censor variable for populations that had not gone extinct by the end of the experiment. Note that an extinction event here was defined as cell numbers of a population declining below the detection limit of the flow cytometer. We treat extinction as an event occurring on the replicate level in each individual treatment.

### Analysis of short-and long-term responses to changes in salinity, in stable and fluctuating temperatures

We analysed the growth of the surviving replicates as assayed at the end of the experiment in the reciprocal transplants using analyses of variance within a mixed effects model (package nlme, version 3.1-131). Growth relative to growth of the evolved control at 22°C in monoculture was the response variable. This normalisation by growth under standard laboratory conditions allows us to correct for evolution occurring merely due to selection for laboratory conditions and further creates a baseline for easy comparison of the selection temperatures in relation to each other. We fitted the following fixed factors in the global model: species (*Chlamydomonas* or *Ostreococcus*), biotic regime (invading or colonising), selection temperature (22°C,26°C,32°C, or fluctuating), and response type (‘short’ for growth rates in the novel salinity after two weeks of culturing in the novel salinity, ‘long’ for growth rates in the novel salinity after evolution in the novel salinity, ‘back’ for growth rates in the ancestral salinity after evolution in the novel salinity). The ancestral and selection salinities can be inferred from the species and biotic regime factors, and therefore selection salinity was not added as an explicit factor. Replicates (Supporting Table 1 for number of surviving replicates in each unique treatment) nested within ‘unique treatment’ were used as random factors. The nesting was necessary as all replicates originally came from the same starting culture, i.e. replicate 1 of any given treatment was not more or less related to replicate 1 in another treatment than it was to, e.g., replicate 5. We ran the model only on samples where the assay temperature was identical to the selection temperature. We started the model with the fixed factors in full interaction, and searched for the model with the lowest AICc scores through the ‘dredge’ function within the MuMIn package (version 1.40.4). The model with the lowest AICc was consequently used. In all cases of model selection by AICc, we used a delta value of > 2 to confirm the best model (Supporting Tables 5 and 6).

### Analysis of local adaptation to temperature

To specifically test whether samples had locally adapted to their selection temperature without risking over parameterising the mixed model, we built a separate mixed effects model using data where the assay salinity was the same as the selection salinity, thus focusing on the temperature dependence of growth rates in the samples’ selection salinity. We used growth rates relative to the evolved control in mono-culture at 22°C as the response variable, and species (*Chlamydomonas* or *Ostreococcus*), biotic regime (invading or colonising), selection temperature (22°C,26°C,32°C, or fluctuating), and assay temperature (22°C,26°C,32°C, or fluctuating) as fixed factors. The random factors and model fitting were as described above (Supporting Tables 7 and 8).

### Analysis of growth rates in decomposed samples

To analyse whether invaders and residents developed a dependence on each other in the short-term, we compared growth rates of the decomposed samples (decomposition after two weeks of co-culture) to growth rates of the same species at the same temperature and salinity in mono-culture *via* a t-test (Supporting Table 15). For the long term-responses, we analysed the decomposed samples (measured at selection temperature) by fitting a mixed model, using the ratio between growth rates of either species after decomposition and growth rates of the species in co-culture as the response variable. We fitted species (*Chlamydomonas* or *Ostreococcus)*, biotic interaction during selection(resident or invader) and selection temperature as the fixed effects. Random effects and model fitting were as described above (Supporting Tables 11 and 12).

### Phenotypic characterisation

In order to estimate the effect of the selection regimes (biotic scenarios, temperature, and salinity) on cell size and total biomass we fitted a mixed model with the full interaction of the parameters species (*Chlamydomonas* or *Ostreococcus*), assay salinity (‘home’ for the assay salinity being equal to the focal species’ *selection* salinity, and ‘away’ for assay salinity being different from the focal species’ selection salinity), biotic regime (invading or colonising) and selection temperature (22°C, 26°C, 32°C, fluctuating). Model fitting and selection proceeded as described above (Supporting Tables 9 and 10 for size, Supporting Tables 16 and 17 for biomass). For the analysis of rates of net primary production in the evolved and decomposed samples specifically, a mixed model was fitted using species (*Chlamydomonas* or *Ostreococcus*), ‘previous interaction’ (invader or resident), assay salinity (including a unique identifier for each salinity in interaction with whether the sample had been decomposed and at which point in time – after 2 weeks, or at the end of the experiment – it had been decomposed, Fig. 1D for an example), selection temperature (22°C, 26°C, 32°C, fluctuating), ‘biotic selection regime’ (colonisers or invaders) as the fixed effects. Model fitting and selection then proceeded as described above (Supporting Tables 13 and 14).

