## Supplementary material for "Presence of a resident species aids invader evolution": All Supporting Information

**Author affiliations:**
[1] Department of Biology, University of Toronto at Mississauga, 3359 Mississauga Road, William G. Davis Building, Mississauga, ON, L5L 1C6, Canada
[2] Station d'Ecologie Théorique et Expérimentale, UMR 5321 Université Paul Sabatier, Moulis, 09200, France
[3] School of Biological Sciences, University of Reading, Reading, UK
[4] Institute of Marine Ecosystem and Fishery Science, Centre for Earth System Research and Sustainability, Universität Hamburg, Olbersweg 24, 22767 Hamburg

### FIGURES

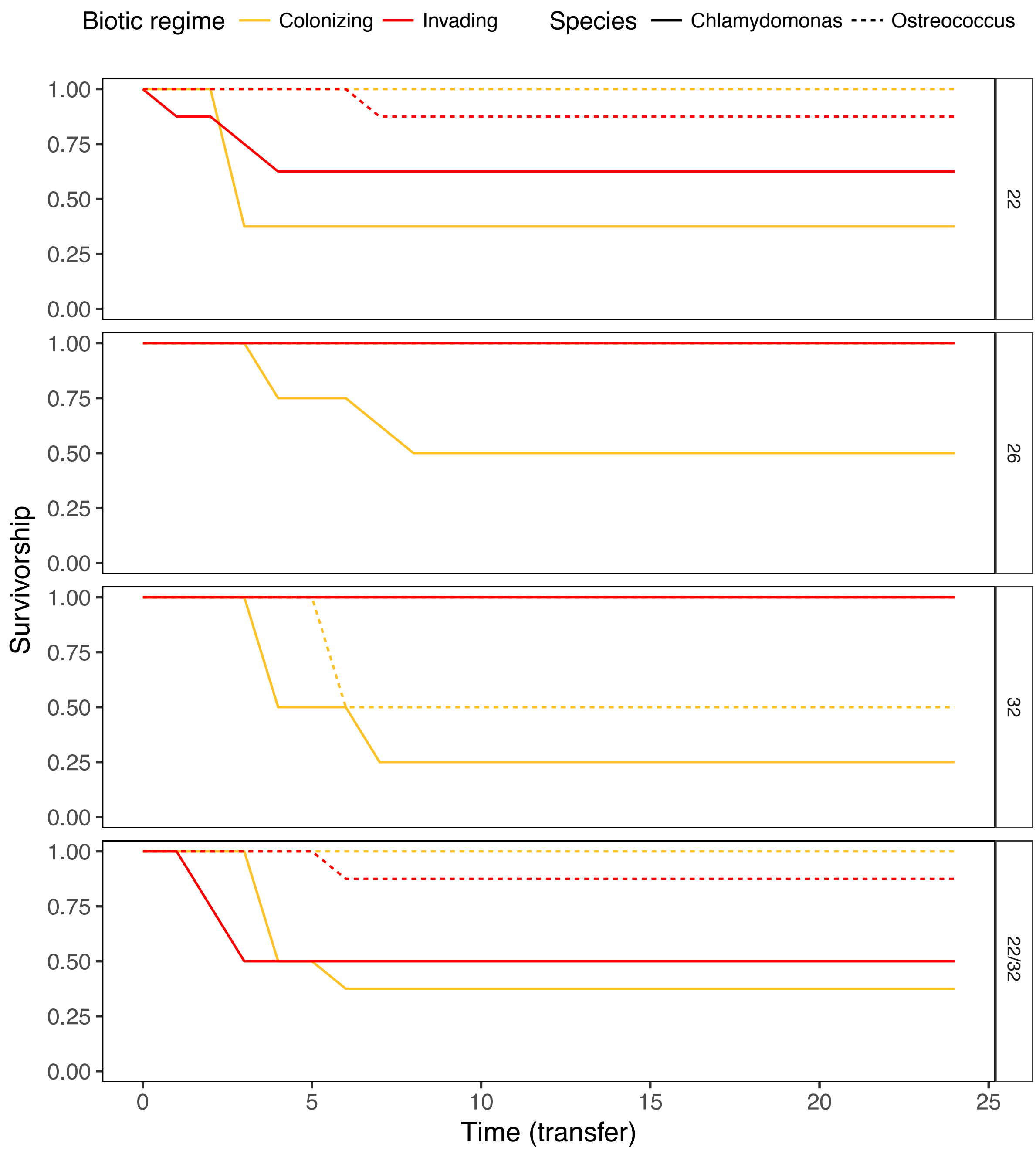

Supporting Figure 1| Survival after transfer into a novel salinity for invaders and colonisers. Survival when invading into a new salinity from rare is enhanced in the presence of a resident species across all selection regimes**.** Displayed is the survivorship for biological replicates of *Chlamydomonas* (solid line) and *Ostreococcus* (dashed line) over time (where one transfer corresponds to one week) and per temperature selection regime, with 1.0 as 100% of populations surviving. Generally, survivorship is enhanced for invaders where there is a resident species present (red) compared to colonisers moving into a novel salinity on their own (orange). This is particularly pronounced under mild warming (26ºC, +4ºC from ambient), where all species invading into an environment where the resident population was present survived. The freshwater species *Chlamydomonas* invading a saltwater environment experienced significantly greater extinction rates at the beginning and in subsequent cycles than the marine species *Ostreococcus* invading a freshwater environment in all temperature regimes.

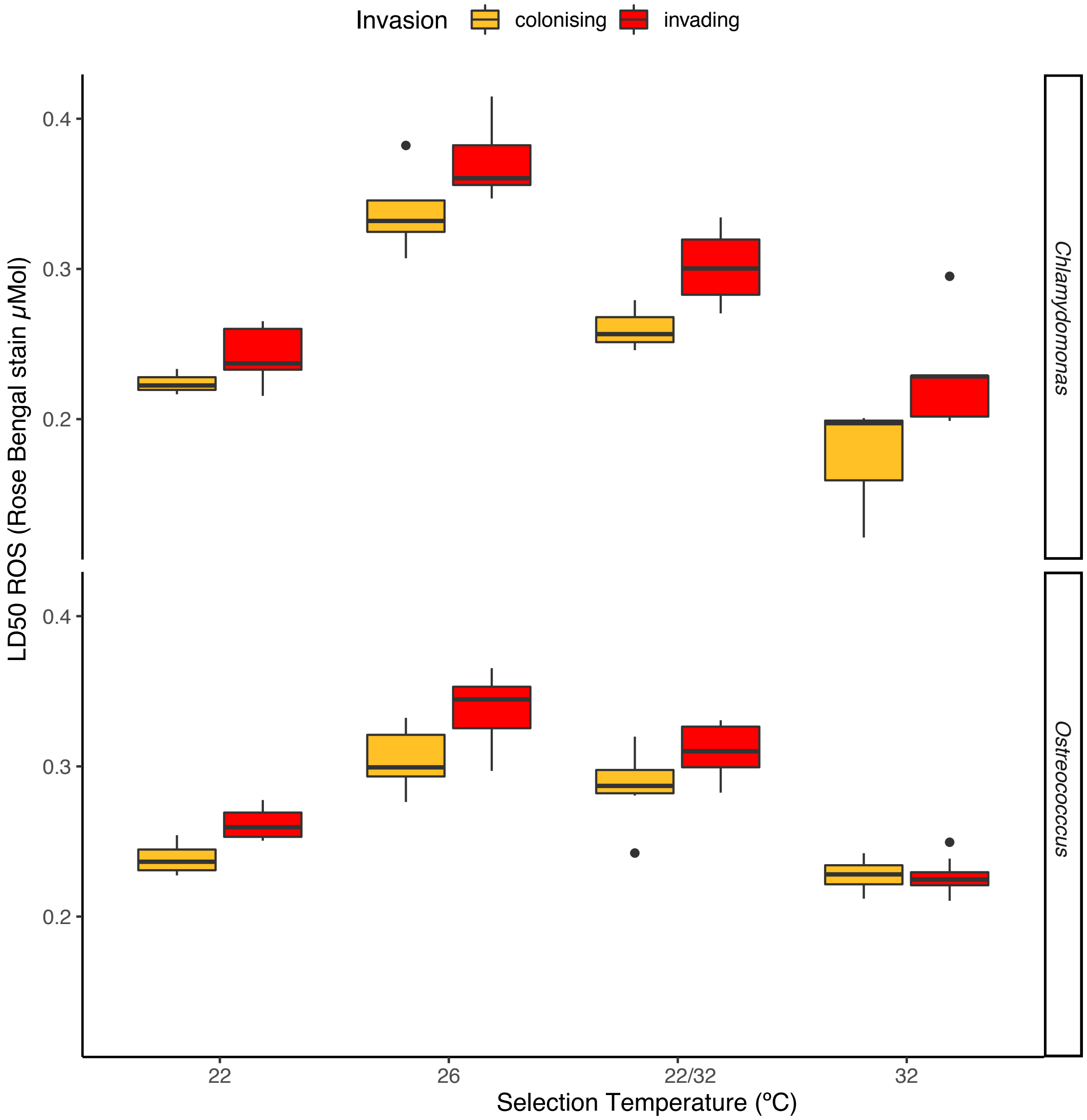

Supporting Figure 2| $LD_{50}$ (in µ Moles) for Rose Bengal stain as a proxy for ROS tolerance in invaders and colonisers across selection temperatures.

ROS tolerance (as per LD50) was highest in the favourable 26ºC and 22/32ºC environments for both species. Across environments, invaders (red) had higher tolerance than colonisers (orange). Boxplots are displayed as is standard, with the belt indicating the median.

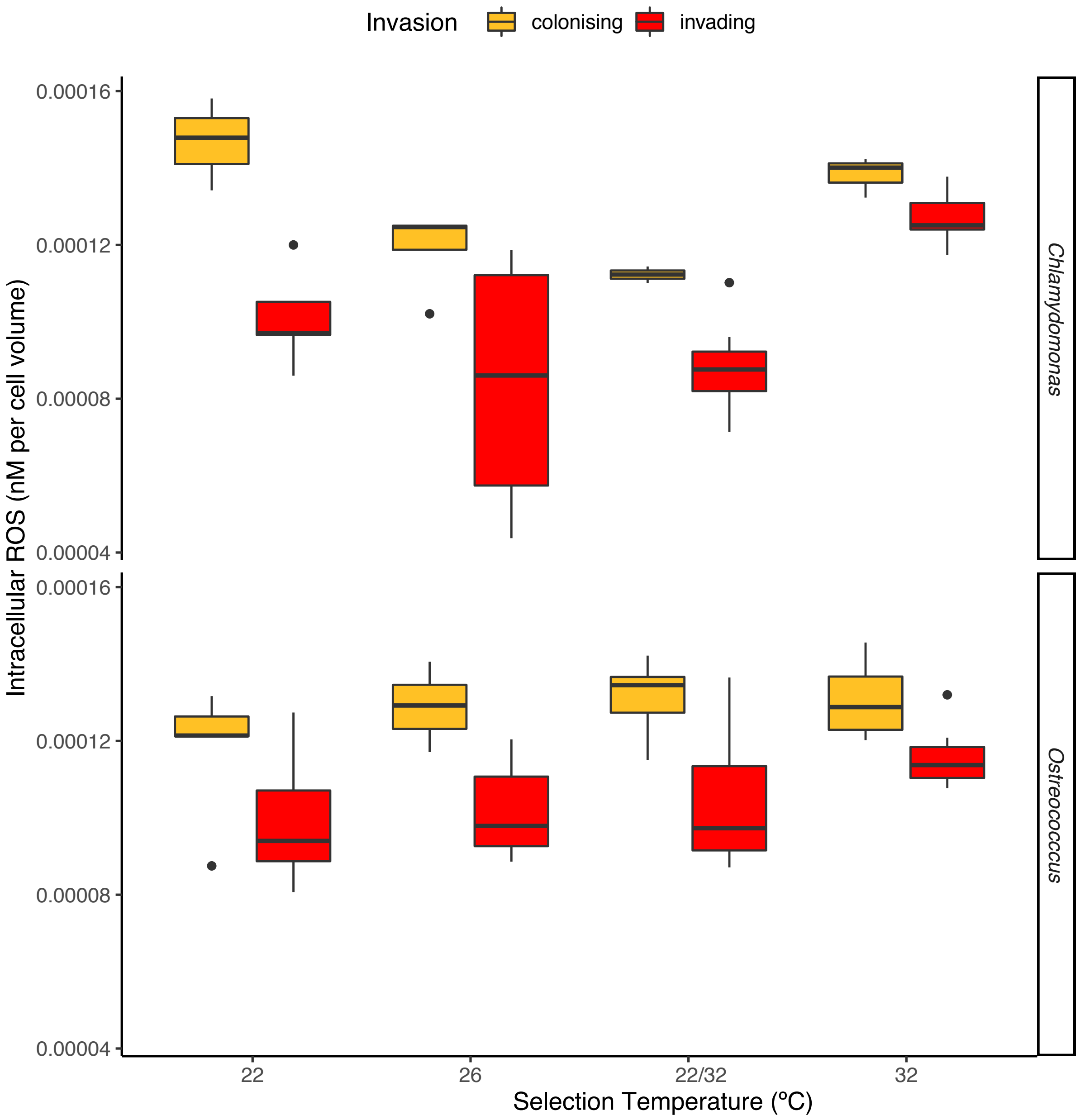

Supporting Figure 3| Intracellular ROS content (corrected for biovolume) in nM established via a fluorescent dichloro-fluorescein stain (2′,7′ dichlorodihydrofluorescein diacetate).

Intracellular ROS was not significantly altered by temperature in *Ostreococccus*, but was lowest in the favourable environments (26ºC and 22/32ºC) for *Chlamydomonas*. Across environments, invaders (red) had lower ROS content than colonisers (orange). Boxplots are displayed as is standard, with the belt indicating the median.

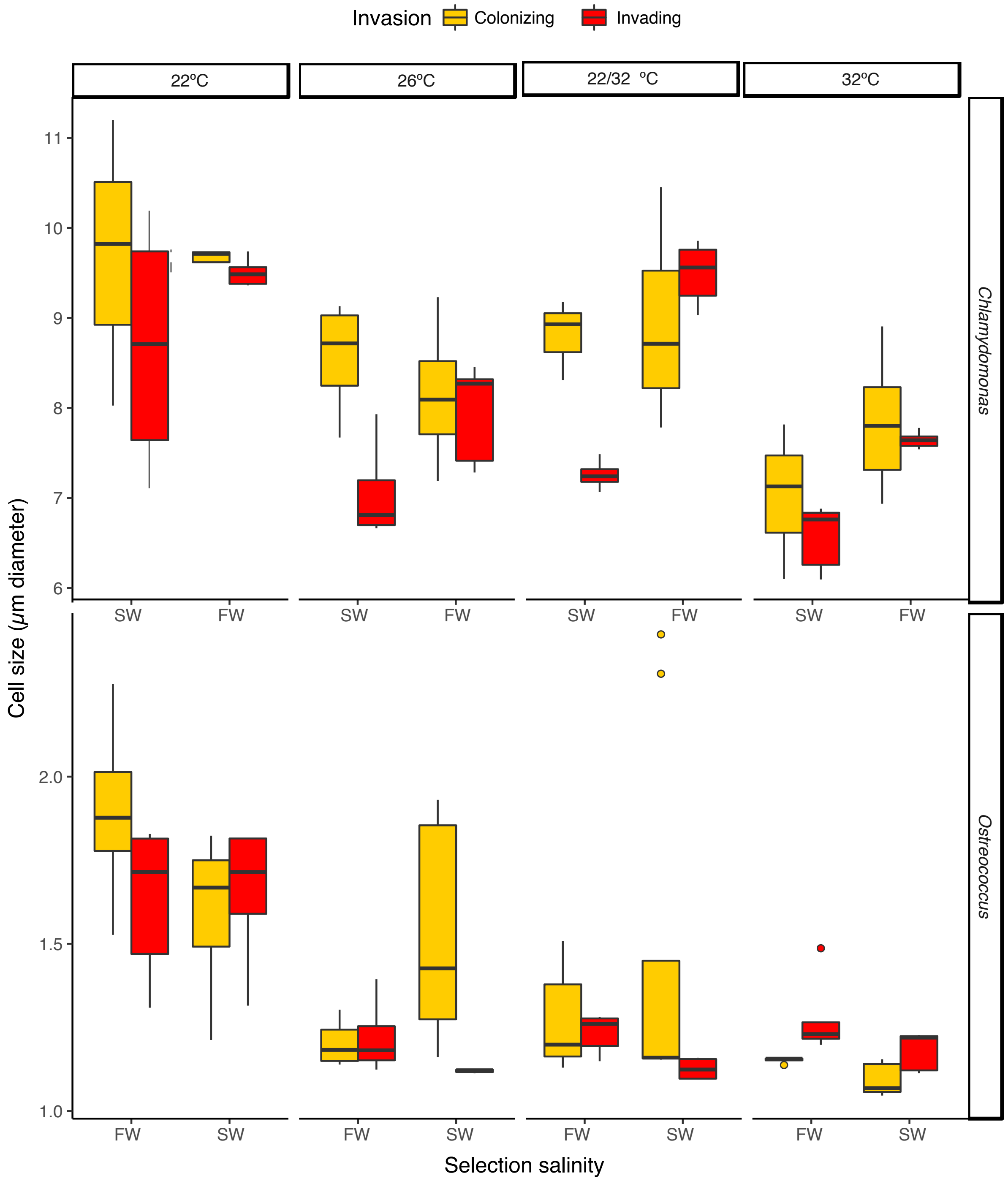

Supporting Figure 4| Size of the focal species is affected by salinity, temperature, and species interactions, but not by salinity alone. We are displaying cell diameters in µm for *Chlamydomonas* (upper panels) and *Ostreococcus* (lower panels) in freshwater (FW) and saltwater (SW) as the salinity that the species were selected in, at all selection temperatures: 22ºC, 26ºC, 32ºC and the fluctuating regime 22/32ºC. Colonisers (orange) are species evolving on their own in either saltwater (SW) or freshwater (FW), and invaders (red) are species evolving with the other species. Warming generally leads to a decrease in cell size, and invaders tend to be smaller than colonisers. Each panel is for one selection temperature and focal species. Boxplots are displayed as is standard, with the belt indicating the median.

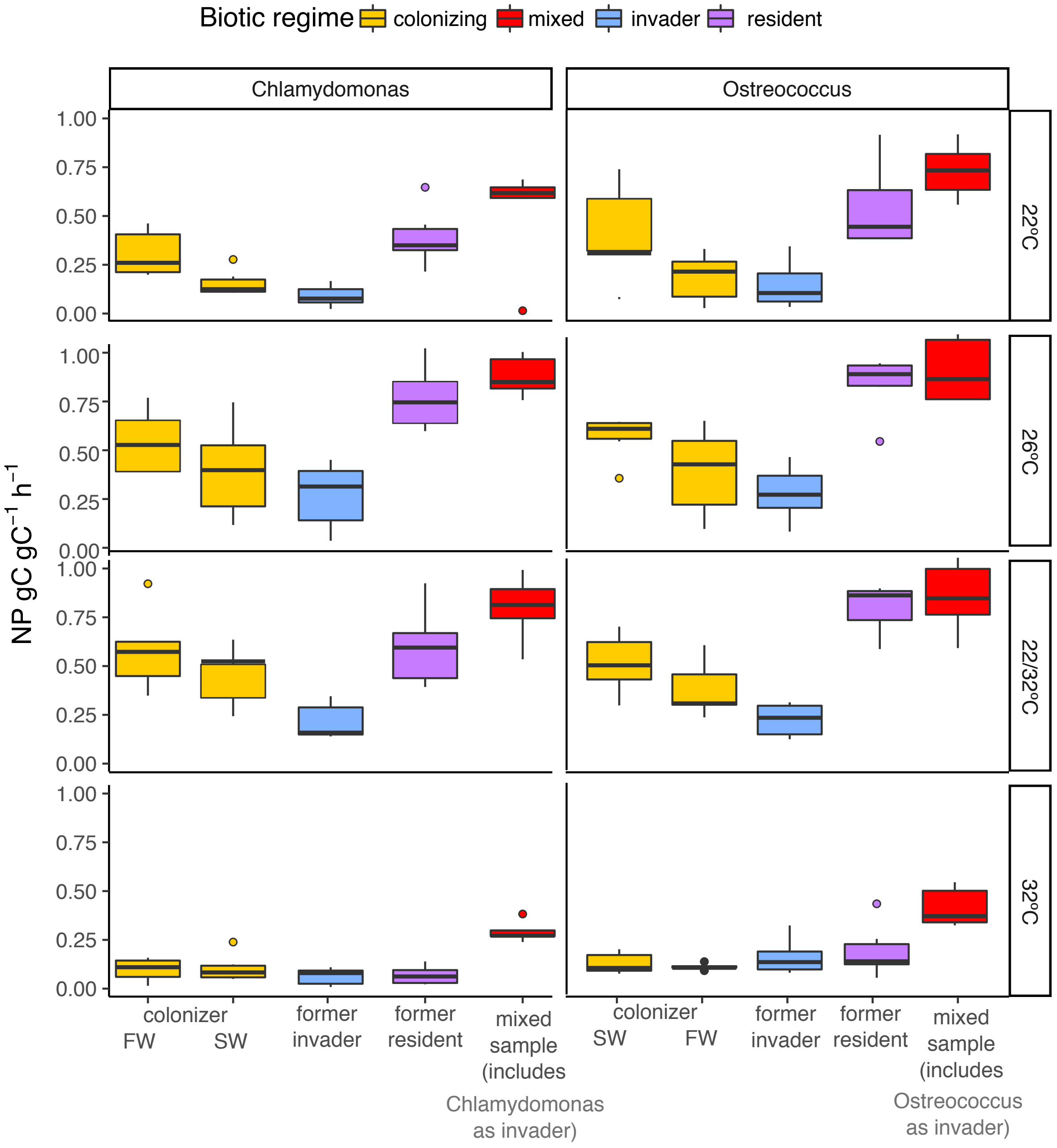

Supporting Figure 5| Net photosynthesis of all evolved and decomposed (after 200 generations) samples at all selection temperatures. Displayed is net primary production as gC per gC per hour. Colonisers (yellow) photosynthesised more when they had evolved in their former home salinity, and net photosynthesis per gC of the mixed community (red) was higher than that of single species isolates. In samples that were deconstructed following 200 generations of selection in a mixed sample, former invaders (blue) always photosynthesised less than the same species after evolution in mono-culture (yellow) -in line with reduced growth rates in these same samples. Former resident species (purple) had higher net photosynthesis rates than the same species after evolution in mono-culture, but only for *Chlamydomonas* did this translate into higher growth rates (Supporting Figure 6). Boxplots are displayed as is standard with the belt indicating the median. n per treatment differed due to differences in extinctions at different selection salinities

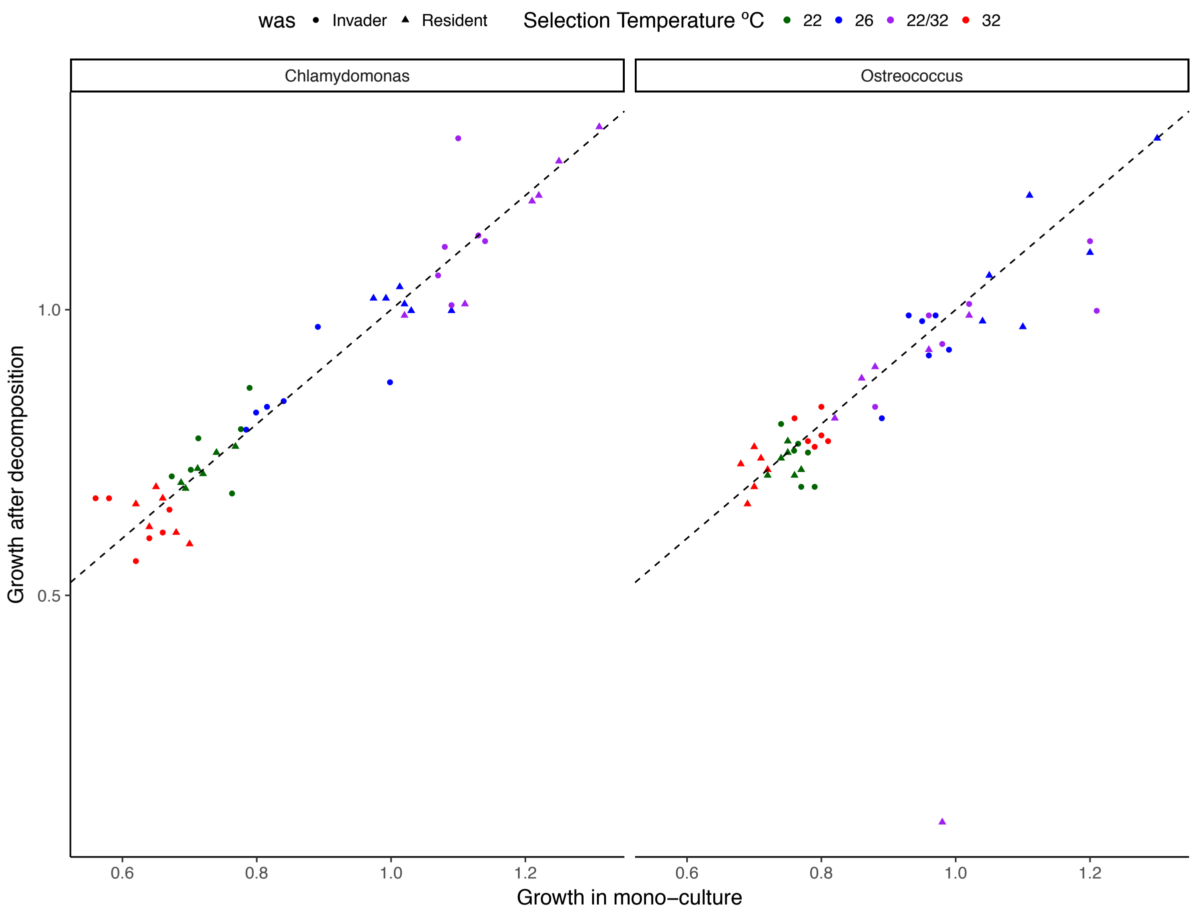

Supporting Figure 6| Comparison of growth rates of invaders and resident species in mono-culture and after decomposition following two weeks of co-culture.

In all selection temperatures and both species, decomposition after two weeks of co-culture did not change growth rates markedly compared to the same species grown in in mono-culture. The dotted line is the 1:1 line, and individual data-points are biological replicates. Colours denote the 22ºC treatment in green, 26ºC in blue, 22/32ºC in purple, and 32ºC in red. In the decomposed samples, triangles indicate former residents, and circles, former invaders.

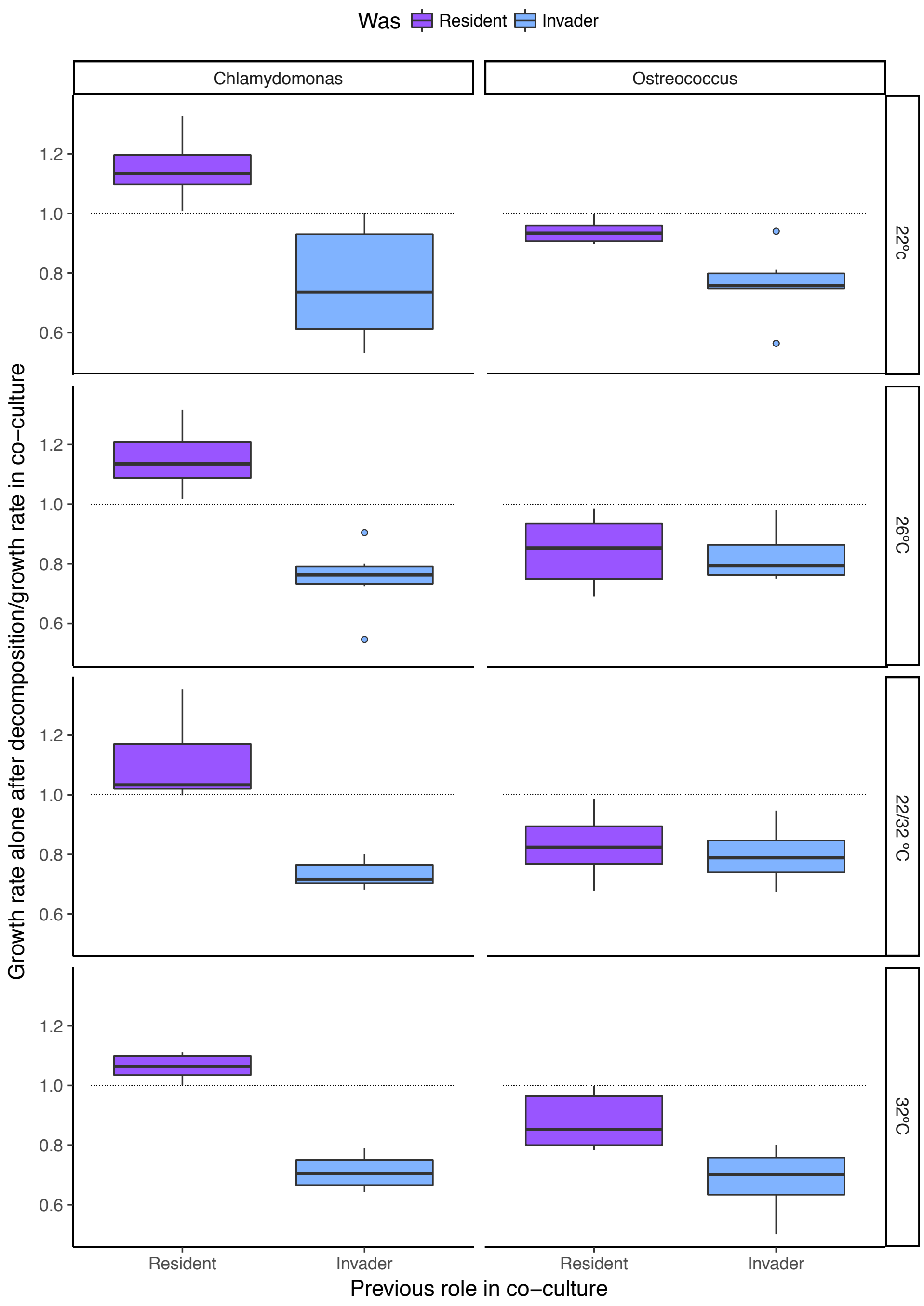

Supporting Figure 7| Growth rates individual species of formerly co-cultured samples after decomposition of the co-cultures. Displayed is the fold change difference in growth rate for former residents (purple) and invaders (blue) after decomposition and growth in co-culture for *Chlamydomonas* (left panel) and *Ostreococcus* (right panel) as the focal species, for all selection temperatures (22ºC, 26ºC, 32ºC and 22/32ºC) at the selection salinity of the mixed sample. The dotted line indicates no change (fold change of 1), values below the line indicate samples growing less well, and values above 1 denote samples growing better than the species in co-culture. Former invaders always grew more poorly upon decomposition, but for former residents, there was a marked species-specific effect: resident *Chlamydomonas* grew better when *Ostreococcus* was removed, whereas resident *Ostreococcus* grew less well upon removal of the invader. See Table S9 and S10 for statistical analysis. Boxplots are displayed as is standard, with the belt indicating the median.

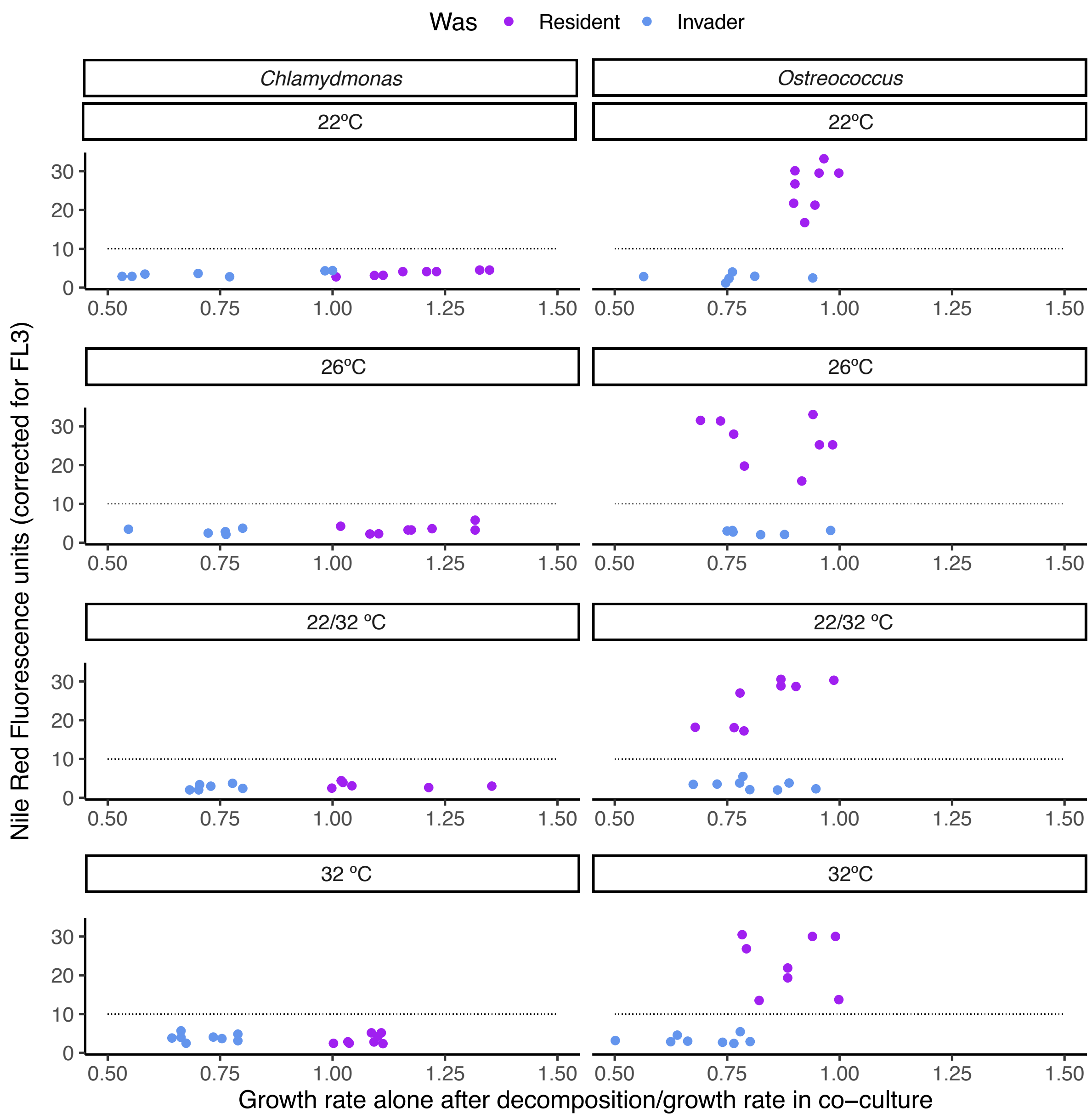

Supporting Figure 8| In samples where surplus NP is not channelled towards growth, cells exhibit elevated Nile Red fluorescence, indicative of lipid storage. As Nile Red excites on the same wavelength as chlorophyll (within the BD Accuri flowcytometer's FL3 channel), all values have been corrected for chlorophyll FL3 in the same sample. Across all temperatures, no Nile Red signal was detectable for the *Chlamydomonas* samples. In *Ostreococcus*, former resident species (purple), which had shown increased NP but no increased growth upon separation from the co-culture, the Nile Red signal was elevated. This was not the case for former *Ostreococcus* invaders (blue). The dotted line at 10 units of fluorescence indicates the detection limit below which we cannot distinguish between background noise and signal. Number of replicates varies due to treatment specific differences in extinction probability.

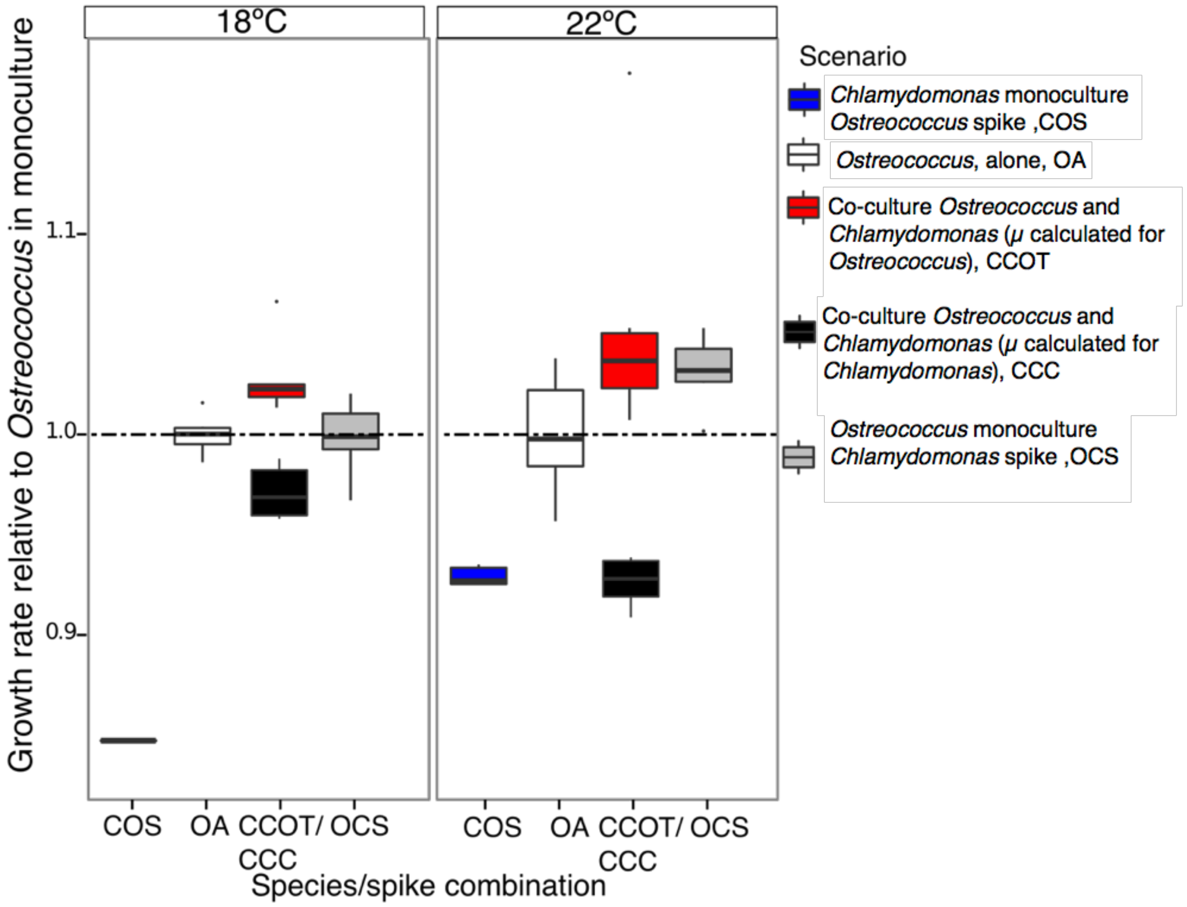

Supporting Figure 9| Result of pilot experiment of Chlamydomonas and Ostreococcus growing in the apparent presence of each other. We measured growth rates of salt-adapted *Chlamydomonas* and *Ostreococcus* at 18ºC and 22ºC in mono-culture (*Ostreococcus:* white), in co-culture (relative growth rate for *Ostreococcus* in red, and for *Chlamydomonas*, in black), as well as in 'spiked' cultures, where 200µl of 0.2 µm filtered dense culture (which allows for molecules, but not organisms to pass) of the marine species where added to the salt-adapted fresh water species and vice versa (*Ostreococcus* spike in *Chlamydomonas* culture: blue, *Chlamydomonas* spike in *Ostreococcus* culture, grey). We also added 200µl of filtered medium from the other species to a 'medium control' but found no effect of 'medium control' on the resident species. Growth rates for all scenarios are displayed as compared to the mean growth rate of *Ostreococcus* in monoculture in the respective incubator. *Ostreococcus* increased its growth rates both in response to a spike from and co-culture with *Chlamydomonas*, and this was particularly pronounced in the 22ºC incubator. Growth rates in *Chlamydomonas*, on the other hand, were reduced both by spiking the sample with *Ostreococcu*s filtrate, and by growing them in co-culture with *Ostreococcus.* This was even more so the case in the 18ºC incubator, were *Chlamydomanos* reacted to the *Ostreococcus* spike by growing substantially more slowly. Additionally, by observing the cells under a microscope, we found that in the co-cultures, there was often had a clear "halo" around single *Chlamydomonas* cells, with *Ostreococcus* aggregating around the outer edge of the halo. Notably, in contrast to our main study, the *Chlamydomonas* strain used in the pilot had already been experimentally adapted to marine conditions, and due to logistic restrictions the pilot had to be carried out under lower light intensities than the main study. n=8 for each individual treatment.

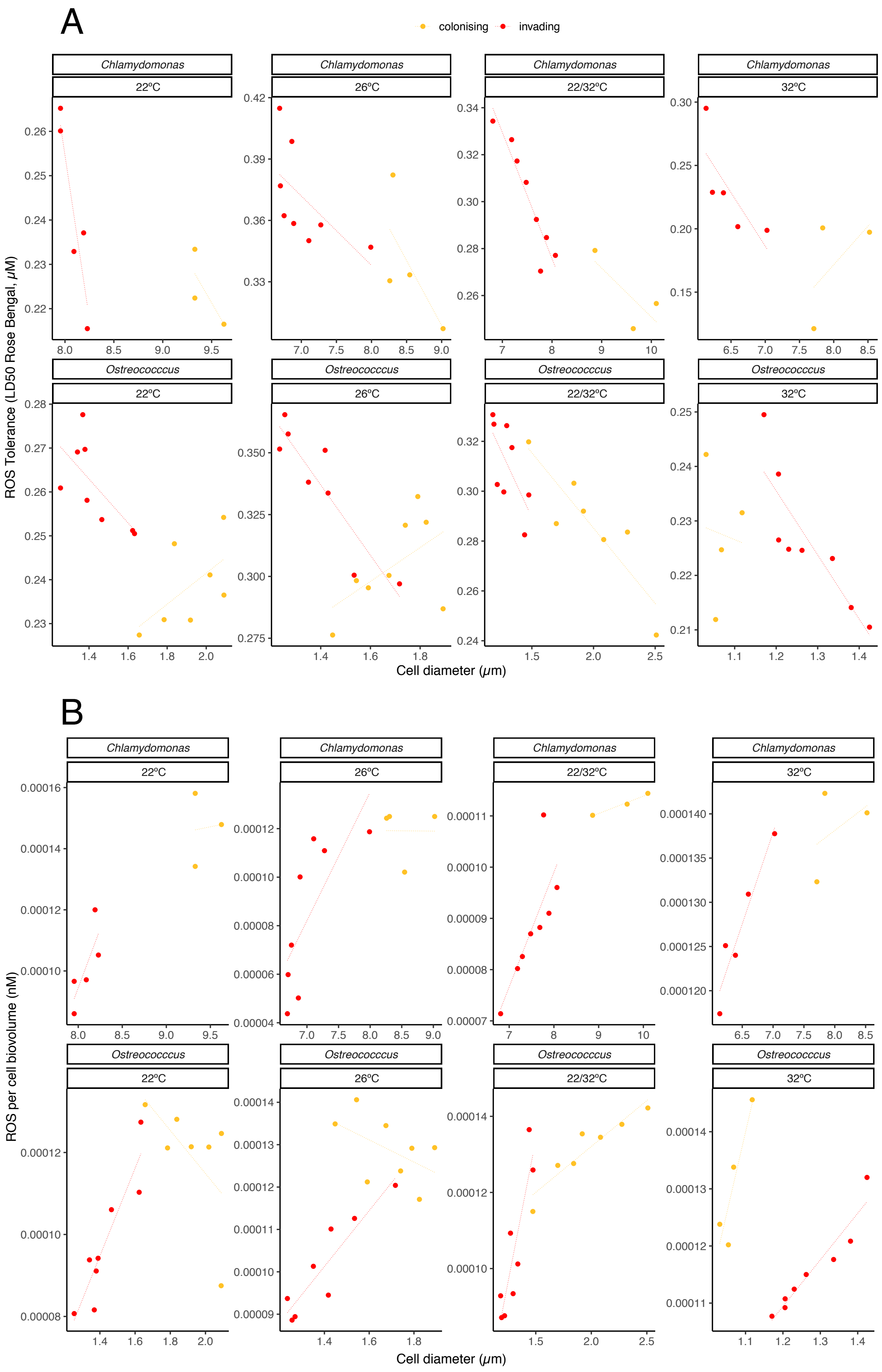
A
colonising
invading
Chlamydomonas
Ostreococcus
22°C
26°C
22/32°C
32°C
ROS Tolerance (LD50 Rose Bengal, μM)
Cell diameter (μm)
B
ROS per cell biovolume (nM)

Supporting Figure 10| Relationship between (A) cell size and $LD_{50}$ (in µ Moles) for Rose Bengal stain as a proxy for ROS tolerance in invaders and colonisers across selection temperatures and (B) cell size and Intracellular ROS content (corrected for biovolume) in nM established via a fluorescent dichloro-fluorescein stain (2′,7′ dichlorodihydrofluorescein diacetate).

Across all temperatures and for both species, larger cells had lower ROS tolerance (lower $LD_{50}$). As invaders (red) tended to be smaller than colonisers (orange), this resulted in higher ROS tolerance for the invaders. Similarly, biovolume-corrected intracellular ROS content was lower in invaders than it was in invaders than colonisers. Lower ROS content in invaders was found even in the case of *Ostreococcus* evolved at 32ºC , where invaders were slightly larger than colonisers (but cells were overall smaller). Number of replicates varies due to treatment specific differences in extinction probability.

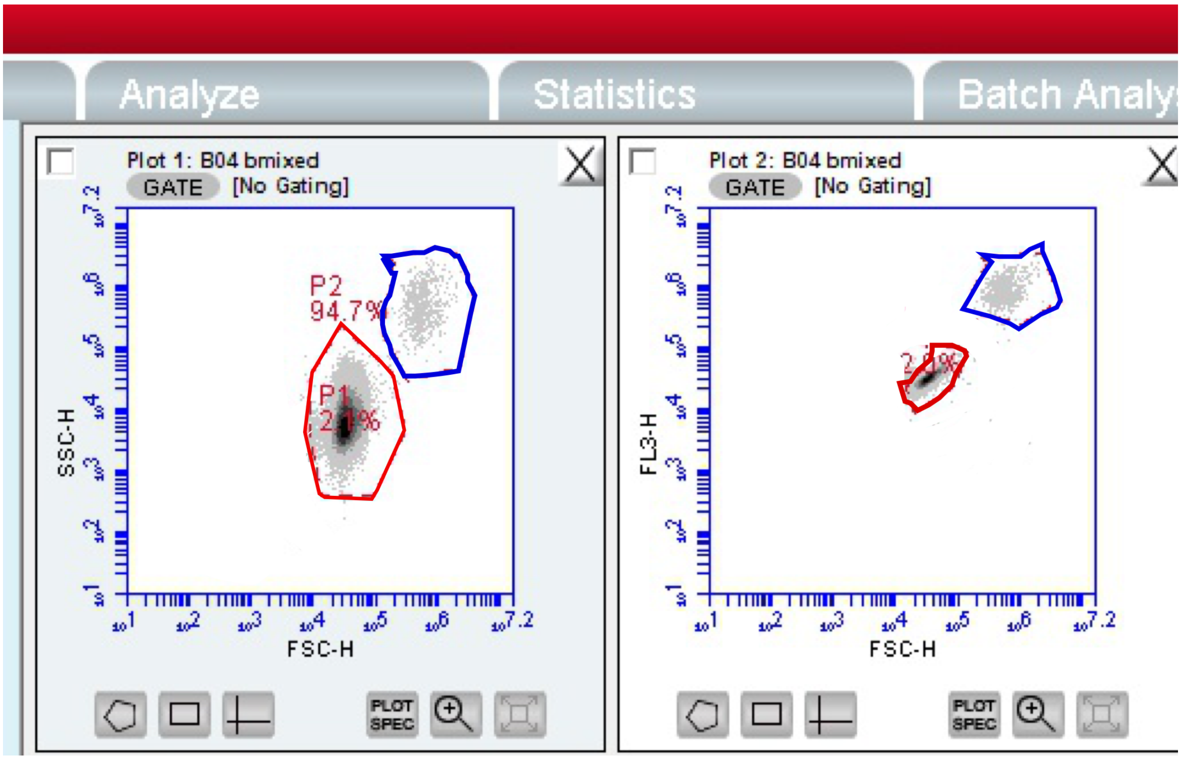

Supporting Figure 11| Screenshot of mixed Chlamydomonas and Ostreococcus population as visualised on the accuri C6 (BD Scientific) flow cytometer used throughout the study. Left: SSC-H (side scatter, proxy for granularity) and FSC-H (forward scatter, proxy for size) for *Ostreococcus* (gated in red) and *Chlamydomonas* (gated in blue). Right: FL3-H (red fluorescence, proxy for chlorophyll content) against FSC-H for *Ostreococcus* and *Chlamydomonas*. The two species can be easily distinguished based on these three parameters alone, which were used in subsequent analyses to determine the number of cells of each species in all treatments at each transfer.

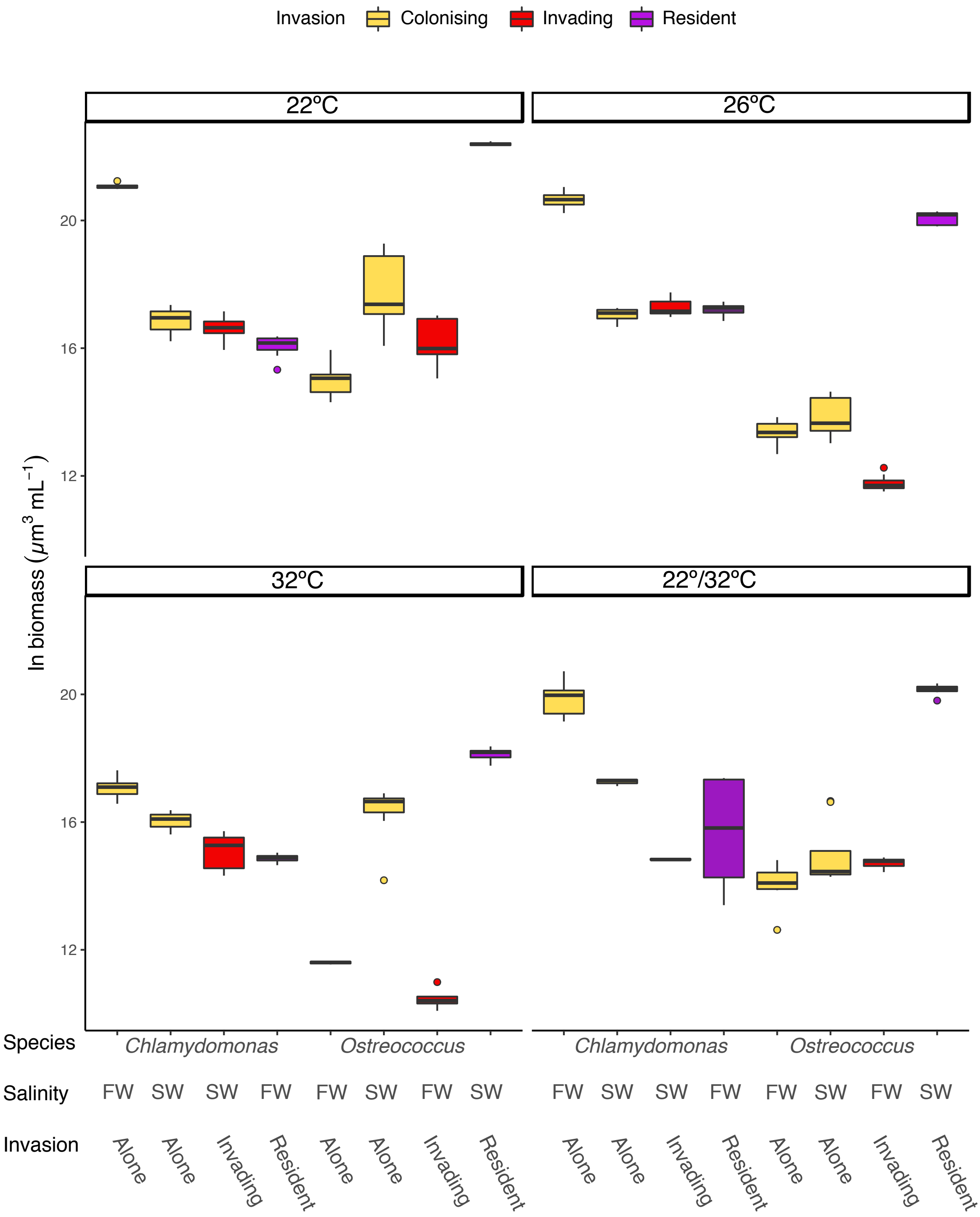

Supporting Figure 12| Total biomass in invaded and invading samples, as well as in monocultures in each selection temperature**.** Biomass in the surviving samples at the end of the experiment differed between selection temperatures as well as biotic scenarios. The lowest biomass was found at 32ºC. When the resident species was invaded by another species, resident biomass was slightly lower than when the resident was growing in its ancestral salinity alone in *Chlamydomonas*, but slightly elevated in *Ostreococcus*. Panels are for the different selection temperatures, and colours indicate the invasion scenarios, with white boxplots for species evolving as colonisers on their own, and grey, as invaders or residents (‘invaded’) in co-culture. FW for freshwater, and SW for saltwater. Boxplots are displayed as is standard with the belt indicating the median. n per treatment differed due to differences in extinctions at different selection salinities (consult Table S2 for details).

### SUPPORTING TABLES

Supporting Table 1| Number of experimental lines surviving by the end of the evolution experiment**.** Each treatment combination had 8 replicate lines at the start of the experiment.

| Focal species | Selection salinity | Biotic scenario | Temperature regime | Number of lines surviving |
|---|---|---|---|---|
| *Chlamydomonas* | freshwater | monoculture | 22 | 8 |
| | | | 26 | 8 |
| | | | 32 | 8 |
| | | | 22/32 | 8 |
| | | coculture (resident) | 22 | 8 |
| | | | 26 | 8 |
| | | | 32 | 8 |
| | | | 22/32 | 8 |
| | saltwater | monoculture | 22 | 3 |
| | | | 26 | 4 |
| | | | 32 | 3 |
| | | | 22/32 | 3 |
| | | coculture (invader) | 22 | 5 |
| | | | 26 | 8 |
| | | | 32 | 8 |
| | | | 22/32 | 4 |
| *Ostreococcus* | freshwater | monoculture | 22 | 8 |
| | | | 26 | 8 |
| | | | 32 | 4 |
| | | | 22/32 | 7 |
| | | coculture (invader) | 22 | 7 |
| | | | 26 | 8 |
| | | | 32 | 8 |
| | | | 22/32 | 8 |
| | saltwater | monoculture | 22 | 8 |
| | | | 26 | 8 |
| | | | 32 | 8 |
| | | | 22/32 | 8 |
| | | coculture (resident) | 22 | 8 |
| | | | 26 | 8 |
| | | | 32 | 8 |
| | | | 22/32 | 8 |

Supporting Table 2| Model output for analysis of survival/extinction. Rates of extinction in a new salinity vary between species and depend on biotic scenario, and temperature regime. Summary statistics table from a survival analysis using a Cox proportional hazards regression model. Abbreviations are as follows: Exp(coef) is the exponential of the coefficient, SE(coef) is the standard error of the coefficient, and z is the Wald statistic calculated as the ratio of the regression coefficient and the standard error. The coefficients represent the difference between groups measured on the log-hazard scale. The exponentiated coefficients are relative hazards. * for p <0.05, ** for p<0.01, *** for p <0.001

| Comparison | Coefficient | Exp(coef) | SE(coef) | z | P value |
|---|---|---|---|---|---|
| Colonising vs. invading | -1.14 | 0.318 | 0.395 | -2.90 | 0.0037 ** |
| 22 vs 26°C | -1.22 | 0.295 | 0.603 | -2.03 | 0.043 * |
| 22 vs 32°C | -0.164 | 0.849 | 0.462 | -0.36 | 0.72 |
| 22 vs 22/32 rapid | -0.0462 | 0.955 | 0.462 | -0.10 | 0.92 |
| *Chlamydomonas vs Ostreococcus* | -1.88 | 0.153 | 0.455 | -4.13 | <0.0001 *** |

Supporting Table 3| Model selection for mixed effects model investigating effects of biotic and abiotic treatments on population size after ~ 70 generations This model is restricted to population size data (cells/mL) after ca 70 generations. The model with the lowest AICc (with a ΔAICc >2), included, through model averaging, selection temperature and selection salinity, but not whether a sample was invaded or colonised.

Identifiers are used as follows: 'Salinity' with Freshwater or Saltwater, Invasion with 'invader' or 'coloniser' and temp for selection temperature (22ºC,26ºC,32ºC, or fluctuating). Note that here, we did not discern between the two species in order to avoid overfitting. Df is for degrees of freedom, logLiK, for log likelihood, AICc as akaike information criterion for small sample sizes, delta as the difference in AICc between the best model and the other models, and weight, as the model weight (i.e.exp( -1/2 * delta) divided by the sum of this for all models) .+ denotes that a component was part of the model. We show the first ten models with lowest AICc out of 20 possible models. Models used for averaging are in bold.

| Model | (Intercept) | Salinity | Invasion | temp | Salinity:Invasion | Salinity:temp | Invasion:temp | Salinity:Invasion:temp | df | logLik | AICc | delta | weight |
|---|---|---|---|---|---|---|---|---|---|---|---|---|---|
| **1** | **49925** | **+** | **NA** | **NA** | **NA** | **NA** | **NA** | **NA** | **4** | **-547.27** | **1103.69** | **0.00** | **0.41** |
| **2** | **109885** | **NA** | **NA** | **NA** | **NA** | **NA** | **NA** | **NA** | **3** | **-548.71** | **1104.09** | **0.40** | **0.33** |
| **3** | **61524** | **+** | **NA** | **+** | **NA** | **NA** | **NA** | **NA** | **5** | **-547.21** | **1106.19** | **2.50** | **0.12** |
| 4 | 122935 | NA | + | NA | NA | NA | NA | NA | 4 | -548.67 | 1106.49 | 2.80 | 0.10 |
| 5 | 68095 | + | + | NA | + | NA | NA | NA | 6 | -547.19 | 1108.93 | 5.24 | 0.03 |
| 6 | 185371 | NA | NA | + | NA | NA | NA | NA | 7 | -547.72 | 1112.94 | 9.25 | 0.00 |
| 7 | 125605 | + | NA | + | NA | NA | NA | NA | 8 | -546.28 | 1113.20 | 9.51 | 0.00 |
| 8 | 198615 | NA | + | + | NA | NA | NA | NA | 8 | -547.68 | 1116.01 | 12.32 | 0.00 |
| 9 | 137205 | + | + | + | NA | NA | NA | NA | 9 | -546.21 | 1116.43 | 12.74 | 0.00 |
| 10 | 143775 | + | + | + | + | NA | NA | NA | 10 | -546.19 | 1119.97 | 16.29 | 0.00 |

Supporting Table 4| Model output on analysis of mixed effects model identified from Supporting Table 3 (population size after ~ 70 generations)

'Df' is degrees of freedom, t-values are the ratios of parameter estimates and standard errors. CI are the upper and lower confidence intervals (95%). Regime identifiers are as above.

| **Regime** | **Estimate** | **CI (lower)** | **CI (upper)** | **Std.Error** | **DF** | **t-value** | **p-value** |
|---|---|---|---|---|---|---|---|
| (Intercept) | 127546.58 | -116306.19 | 371399.34 | 11862.58 | 26 | 1.0751 | 0.2920 |
| temp: 22/32 | -87055.71 | -428653.13 | 254541.72 | 16614.64 | 26 | -0.5238 | 0.6050 |
| temp: 26 | -90086.63 | -431684.06 | 251510.79 | 16684.64 | 26 | -0.5421 | 0.5920 |
| temp: 32 | -124587.32 | -466184.75 | 217010.11 | 16684.64 | 26 | -0.7497 | 0.4600 |
| Salinity: SW | 118255.13 | -347554.08 | 584064.34 | 16771.81 | 4 | 0.7049 | 0.5200 |
| temp22/32: SalinitySW | 22277.89 | -460813.82 | 505369.60 | 23520.58 | 26 | 0.0948 | 0.9250 |
| temp26: SalinitySW | 64084.01 | -419007.7 | 547175.72 | 23520.58 | 26 | 0.2727 | 0.7870 |
| temp32: SalinitySW | 36528.76 | -446562.96 | 519620.47 | 23520.58 | 26 | 0.1554 | 0.8780 |

Supporting Table 5| Model selection for mixed effects model investigating the impact of all biotic and abiotic effects on growth rates during the reciprocal transplant experiment: This model is restricted to data from the reciprocal transplant where cultures were assayed in the temperature regime they had evolved in. The full model had the lowest AICc with a ΔAICc >>2, and was therefore used for further analysis. Abbreviations are used as follows: Intc for intercept, rt for 'response type' ('short' for growth rates in the novel salinity after two weeks of culturing in the novel salinity, 'long' for growth rates in the novel salinity after evolution in the novel salinity, 'back' for growth rates in the ancestral salinity after evolution in the novel salinity), Inv for biotic regime (invading or colonising), sT for selection temperature (22ºC,26ºC,32ºC, or fluctuating), and sp for species (*Chlamydomonas* or *Ostreococcus*). Df is for degrees of freedom, logLiK, for log likelihood, AICc as akaike information criterion for small sample sizes, delta as the difference in AICc between the best model and the other models, and weight, as the model weight (i.e.exp( -1/2 * delta) divided by the sum of this for all models) .+ denotes that a component was part of the model. We show the first ten models with lowest AICc out of 167 possible models.

| | **Intc** | **rt** | **Inv** | **sT** | **sp** | **rt: Inv** | **rt: sT** | **rt: sp** | **Inv: sT** | **Inv: sp** | **sT: sp** | **rt:Inv: sT** | **rt:Inv: sp** | **rt:sT: who** | **Inv:sT:sp** | **rt:In: sT:sp** | **df** | **Log Lik** | **AICc** | **delta** | **weight** |
|---|---|---|---|---|---|---|---|---|---|---|---|---|---|---|---|---|---|---|---|---|---|
| 1 | **0.67** | + | + | + | + | + | + | + | + | + | + | + | + | + | + | + | **51** | **243.40** | **-364.71** | **0.00** | **0.99** |
| 2 | 0.69 | + | + | + | + | + | + | + | + | + | + | NA | NA | + | + | NA | 37 | 215.46 | -346.81 | 17.90 | 0.00 |
| 3 | 0.72 | + | + | + | + | + | + | + | + | + | + | + | NA | + | + | NA | 43 | 222.83 | -345.75 | 18.95 | 0.00 |
| 4 | 0.71 | + | + | + | + | + | + | + | + | + | + | NA | + | + | + | NA | 39 | 217.33 | -345.36 | 19.35 | 0.00 |
| 5 | 0.74 | + | + | + | + | + | + | + | + | + | + | + | + | + | + | NA | 45 | 224.76 | -344.19 | 20.52 | 0.00 |
| 6 | 0.71 | + | + | + | + | NA | + | + | + | + | + | NA | NA | + | + | NA | 35 | 201.95 | -324.89 | 39.82 | 0.00 |
| 7 | 0.70 | + | + | + | + | + | NA | NA | + | + | + | NA | NA | NA | + | NA | 23 | 187.02 | -324.26 | 40.45 | 0.00 |
| 8 | 0.67 | + | + | + | + | + | + | NA | + | + | + | NA | NA | NA | + | NA | 29 | 193.51 | -322.94 | 41.77 | 0.00 |
| 9 | 0.71 | + | + | + | + | + | NA | + | + | + | + | NA | NA | NA | + | NA | 25 | 188.42 | -322.37 | 42.34 | 0.00 |
| 10 | 0.70 | + | + | + | + | + | + | NA | + | + | + | + | NA | NA | + | NA | 35 | 200.35 | -321.71 | 43.00 | 0.00 |

Supporting Table 6| Model output on analysis of mixed effects model identified from Supporting Table 5 (effects on growth rates). 'Df' is degrees of freedom, t-values are the ratios of parameter estimates and standard errors. p values <0.05 are in bold. Abbreviations of model parameters are as in Supporting Table 5 above. CI are the upper and lower confidence intervals (95%).

| | **Estimate** | **CI (lower)** | **CI (upper)** | **Std. Error** | **DF** | **t-value** | **p-value** | |
|---|---|---|---|---|---|---|---|---|
| **(Intercept, i.e Chlamydomonas colonising its ancestral environment at 22ºC in the short term)** | 0.67 | 0.57 | 0.76 | 0.05 | 224 | 13.43 | **<0.001** | *** |
| **Inv invading** | 0.27 | 0.14 | 0.40 | 0.06 | 44 | 4.32 | **<0.001** | *** |
| spOstreococcus | -0.10 | -0.23 | 0.04 | 0.07 | 44 | -1.47 | 0.148 | |
| **rt Long** | 0.37 | 0.20 | 0.54 | 0.09 | 224 | 4.26 | **<0.001** | *** |
| **rt Back** | 0.20 | 0.03 | 0.37 | 0.09 | 224 | 2.30 | **0.022** | * |
| **sT 26** | 0.20 | 0.07 | 0.33 | 0.07 | 224 | 3.00 | **0.003** | ** |
| **sT 22/32** | 0.48 | 0.36 | 0.61 | 0.07 | 224 | 7.38 | **<0.001** | *** |
| **sT 32** | 0.17 | 0.03 | 0.30 | 0.07 | 224 | 2.46 | **0.015** | * |
| Invading:spOstreococcus | 0.15 | -0.03 | 0.32 | 0.09 | 44 | 1.68 | 0.101 | |
| Invading: rt Long | -0.12 | -0.33 | 0.10 | 0.11 | 224 | -1.09 | 0.276 | |
| Invading: rt Back | -0.14 | -0.36 | 0.07 | 0.11 | 224 | -1.32 | 0.188 | |
| spOstreococcus: rt Long | -0.06 | -0.27 | 0.14 | 0.11 | 224 | -0.61 | 0.542 | |
| spOstreococcus: rt Back | -0.16 | -0.37 | 0.04 | 0.11 | 224 | -1.56 | 0.12 | |
| Invading: sT 26 | 0.12 | -0.06 | 0.30 | 0.09 | 224 | 1.29 | 0.199 | |
| Invading: sT 22/32 | 0.17 | -0.01 | 0.36 | 0.09 | 224 | 1.86 | 0.064 | . |
| **Invading: sT 32** | -0.24 | -0.42 | -0.05 | 0.09 | 224 | -2.48 | **0.014** | * |
| spOstreococcus: sT 26 | -0.11 | -0.30 | 0.07 | 0.09 | 224 | -1.22 | 0.222 | |
| **spOstreococcus: sT 22/32** | -0.22 | -0.39 | -0.04 | 0.09 | 224 | -2.40 | **0.017** | * |
| **spOstreococcus: sT 32** | -0.34 | -0.52 | -0.16 | 0.09 | 224 | -3.78 | **<0.001** | *** |
| rt Long: sT 26 | 0.16 | -0.06 | 0.38 | 0.11 | 224 | 1.41 | 0.16 | |
| **rt Back: sT 26** | -0.23 | -0.45 | -0.01 | 0.11 | 224 | -2.03 | **0.044** | * |
| rt Long: sT 22/32 | -0.16 | -0.40 | 0.07 | 0.12 | 224 | -1.35 | 0.178 | |
| **rt Back: sT 22/32** | -0.38 | -0.61 | -0.15 | 0.12 | 224 | -3.19 | **0.002** | ** |
| rt Long: sT 32 | -0.21 | -0.44 | 0.03 | 0.12 | 224 | -1.71 | 0.088 | . |
| rt Back: sT 32 | -0.10 | -0.35 | 0.16 | 0.13 | 224 | -0.74 | 0.461 | |
| Invading:spOstreococcus:rt Long | 0.19 | -0.08 | 0.47 | 0.14 | 224 | 1.39 | 0.167 | |
| **Invading:spOstreococcus:rt Back** | 0.43 | 0.16 | 0.71 | 0.14 | 224 | 3.13 | **0.002** | ** |
| Invading:spOstreococcus: sT 26 | 0.18 | -0.07 | 0.44 | 0.13 | 224 | 1.41 | 0.16 | |
| Invading:spOstreococcus: sT 22/32 | -0.11 | -0.36 | 0.14 | 0.13 | 224 | -0.85 | 0.396 | |
| Invading:spOstreococcus: sT 32 | 0.14 | -0.12 | 0.39 | 0.13 | 224 | 1.07 | 0.285 | |
| Invading:rt Long: sT 26 | -0.02 | -0.31 | 0.27 | 0.15 | 224 | -0.13 | 0.895 | |
| **Invading:rt Back: sT 26** | 0.51 | 0.22 | 0.80 | 0.15 | 224 | 3.41 | **0.001** | *** |
| Invading:rt Long: sT 22/32 | 0.03 | -0.28 | 0.33 | 0.15 | 224 | 0.18 | 0.86 | |
| Invading:rt Back: sT 22/32 | 0.25 | -0.05 | 0.55 | 0.15 | 224 | 1.61 | 0.108 | |

| | | | | | | | | |
|---|---|---|---|---|---|---|---|---|
| Invading:rt Long: sT 32 | 0.19 | -0.11 | 0.50 | 0.15 | 224 | 1.25 | 0.212 | |
| Invading:rt Back: sT 32 | 0.07 | -0.24 | 0.39 | 0.16 | 224 | 0.46 | 0.648 | |
| spOstreococcus:rt Long: sT 26 | -0.01 | -0.30 | 0.28 | 0.14 | 224 | -0.07 | 0.946 | |
| spOstreococcus:rt Back: sT 26 | 0.20 | -0.09 | 0.48 | 0.14 | 224 | 1.35 | 0.177 | |
| **spOstreococcus:rt Long: sT 22/32** | 0.38 | 0.09 | 0.67 | 0.15 | 224 | 2.56 | **0.011** | * |
| **spOstreococcus:rt Back: sT 22/32** | 0.56 | 0.27 | 0.85 | 0.15 | 224 | 3.81 | **<0.001** | *** |
| **spOstreococcus:rt Long: sT 32** | 0.46 | 0.16 | 0.76 | 0.15 | 224 | 2.99 | **0.003** | ** |
| spOstreococcus:rt Back: sT 32 | 0.13 | -0.19 | 0.45 | 0.16 | 224 | 0.80 | 0.422 | |
| **Invading:spOstreococcus:Long:sT 26** | -0.46 | -0.84 | -0.08 | 0.19 | 224 | -2.36 | **0.019** | * |
| **Invading:spOstreococcus:Back:sT 26** | -1.01 | -1.39 | -0.63 | 0.19 | 224 | -5.20 | **<0.001** | *** |
| **Invading:spOstreococcus:Long:sT 22/32** | -0.40 | -0.79 | -0.02 | 0.20 | 224 | -2.06 | **0.041** | * |
| **Invading:spOstreococcus:Back:sT 22/32** | -0.72 | -1.11 | -0.34 | 0.20 | 224 | -3.67 | **<0.001** | *** |
| **Invading:spOstreococcus:Long:sT 32** | -0.45 | -0.85 | -0.05 | 0.20 | 224 | -2.24 | **0.026** | * |
| Invading:spOstreococcus:Back:sT 32 | -0.37 | -0.78 | 0.04 | 0.21 | 224 | -1.78 | 0.076 | . |

Supporting Table 7| Model selection table for linear mixed model on local adaptation to temperature. The global model is restricted to data from the reciprocal transplant where samples were assayed in their selection salinity, but in a full reciprocal of all selection temperatures. The best model included the interaction of selection temperature, assay temperature and biotic regime, but not the species. Abbreviations are used as follows: Intc for intercept, sT for selection temperature (22ºC,26ºC, 32ºC or 22/32ºC), aT for assay temperature (22ºC,26ºC, 32ºC or 22/32ºC), Inv for biotic regime (colonising or invading), sp for species (*Ostreococcus* or *Chlamydomonas*). Further, df, for degrees of freedom, logLiK, for log likelihood, AICc as akaike information criterion for small sample sizes, delta as the difference in AICc between the best model and the other models, and weight, as the model weight (i.e. exp( -1/2 * delta) divided by the sum of this for all models) .+ denotes that a component was part of the model. We display the first ten out of 166 possible models

| **Model** | **Intc** | **aT** | **sT** | **sp** | **Inv** | **aT * sT** | **aT* sp** | **aT * Inv** | **sT* sp** | **sT* Inv** | **Sp* Inv** | **aT* sT * sp** | **aT * sT * inv** | **aT * sp * Inv** | **sT* sp * Inv** | **aT * sT * Inv * sp** | **df** | **logLik** | **AICc** | **delta** | **weight** |
|---|---|---|---|---|---|---|---|---|---|---|---|---|---|---|---|---|---|---|---|---|---|
| **1** | **0.95** | **+** | **+** | **+** | **+** | **+** | **+** | **+** | **+** | **+** | **+** | **+** | **+** | **NA** | **+** | **NA** | **55** | **408.83** | **-685.65** | **0.00** | **0.73** |
| 2 | 1.03 | + | + | + | + | + | + | + | + | + | + | + | + | + | + | + | 67 | 424.71 | -683.78 | 4.00 | 0.18 |
| 3 | 0.97 | + | + | + | + | + | + | + | + | + | + | + | + | + | + | NA | 58 | 411.91 | -683.62 | 5.87 | 0.08 |
| 4 | 0.90 | + | + | + | + | + | + | + | + | + | + | + | + | NA | NA | NA | 52 | 403.37 | -678.09 | 6.03 | 0.00 |
| 5 | 0.89 | + | + | + | + | + | + | + | + | + | NA | + | + | NA | NA | NA | 51 | 401.74 | -671.75 | 11.56 | 0.00 |
| 6 | 0.92 | + | + | + | + | + | + | + | + | + | + | + | + | + | NA | NA | 55 | 406.26 | -665.21 | 17.90 | 0.00 |
| 7 | 0.95 | + | + | + | + | + | NA | + | + | + | NA | NA | + | NA | NA | NA | 39 | 364.92 | -643.03 | 24.44 | 0.00 |
| 8 | 1.01 | + | + | + | + | + | + | + | + | + | + | NA | + | NA | + | NA | 46 | 373.68 | -642.93 | 46.62 | 0.00 |
| 9 | 1.03 | + | + | + | + | + | NA | + | + | + | + | NA | + | NA | + | NA | 43 | 369.81 | -642.36 | 46.72 | 0.00 |
| 10 | 0.94 | + | + | + | + | + | + | + | + | + | NA | NA | + | NA | NA | NA | 42 | 368.27 | -685.65 | 47.29 | 0.00 |

Supporting Table 8| Model output on analysis of mixed effects model identified from Supporting Table 7 **(temperature).** 'Df' is degrees of freedom; t-values are the ratios of parameter estimates and standard errors. Abbreviations of model parameters are as in Supporting Table 7 above. CI are the upper and lower confidence intervals (95%).

| | **Estimate** | **CI (lower)** | **CI (upper)** | **Std.Error** | **DF** | **t-value** | **p-value** | |
|---|---|---|---|---|---|---|---|---|
| (Intercept, i.e Chlamydomonas colonising its ancestral environment at 22ºC in the short term) | 0.90 | 0.82 | 0.97 | 0.04 | 273 | 24.94 | <0.001 | *** |
| sT 26 | 0.13 | 0.04 | 0.22 | 0.05 | 62 | 2.77 | 0.007 | ** |
| sT 22/32 | 0.50 | 0.40 | 0.59 | 0.05 | 62 | 10.56 | <0.001 | *** |
| sT 32 | -0.25 | -0.36 | -0.14 | 0.06 | 62 | -4.51 | <0.001 | *** |
| Invasion: invading | 0.43 | 0.34 | 0.52 | 0.05 | 273 | 9.14 | <0.001 | *** |
| aT 26 | 0.35 | 0.27 | 0.43 | 0.04 | 273 | 9.03 | <0.001 | *** |
| aT 22/32 | 0.39 | 0.32 | 0.47 | 0.04 | 273 | 10.08 | <0.001 | *** |
| aT 32 | -0.33 | -0.41 | -0.25 | 0.04 | 273 | -8.48 | <0.001 | *** |
| sT 26:Invading | -0.58 | -0.70 | -0.46 | 0.06 | 62 | -9.57 | <0.001 | *** |
| sT 22/32:Invading | -0.52 | -0.64 | -0.40 | 0.06 | 62 | -8.42 | <0.001 | *** |
| sT 32:Invading | -0.29 | -0.42 | -0.15 | 0.07 | 62 | -4.25 | <0.001 | *** |
| sT 26: aT 26 | -0.18 | -0.29 | -0.08 | 0.05 | 273 | -3.43 | 0.001 | *** |
| sT 22/32: aT 26 | -0.47 | -0.58 | -0.36 | 0.06 | 273 | -8.53 | <0.001 | *** |
| sT 32: aT 26 | -0.37 | -0.49 | -0.24 | 0.06 | 273 | -5.85 | <0.001 | *** |
| sT 26: aT 22/32 | -0.26 | -0.37 | -0.16 | 0.05 | 273 | -4.88 | <0.001 | *** |
| sT 22/32: aT 22/32 | -0.40 | -0.50 | -0.29 | 0.06 | 273 | -7.20 | <0.001 | *** |
| sT 32: aT 22/32 | -0.32 | -0.44 | -0.20 | 0.06 | 273 | -5.15 | <0.001 | *** |
| sT 26: aT 32 | 0.28 | 0.17 | 0.38 | 0.05 | 273 | 5.15 | <0.001 | *** |
| sT 22/32: aT 32 | 0.19 | 0.09 | 0.30 | 0.06 | 273 | 3.52 | 0.001 | *** |
| sT 32: aT 32 | 0.66 | 0.53 | 0.78 | 0.06 | 273 | 10.53 | <0.001 | *** |
| Invading: aT 26 | -0.18 | -0.28 | -0.07 | 0.05 | 273 | -3.37 | 0.001 | *** |
| Invading: aT 22/32 | -0.34 | -0.45 | -0.23 | 0.06 | 273 | -6.17 | <0.001 | *** |

| | | | | | | | | |
|---|---|---|---|---|---|---|---|---|
| Invading: aT 32 | -0.33 | -0.44 | -0.22 | 0.06 | 273 | -5.95 | <0.001 | *** |
| sT 26:Invading:aT 26 | 0.68 | 0.54 | 0.82 | 0.07 | 273 | 9.43 | <0.001 | *** |
| sT 22/32:Invading:aT 26 | 0.41 | 0.27 | 0.56 | 0.07 | 273 | 5.57 | <0.001 | *** |
| sT 32:Invading:aT 26 | 0.18 | 0.02 | 0.33 | 0.08 | 273 | 2.26 | 0.025 | * |
| sT 26:Invading:aT 22/32 | 0.64 | 0.49 | 0.78 | 0.07 | 273 | 8.59 | <0.001 | *** |
| sT 22/32:Invading:aT 22/32 | 0.68 | 0.53 | 0.83 | 0.08 | 273 | 9.01 | <0.001 | *** |
| sT 32:Invading:aT 22/32 | 0.28 | 0.13 | 0.44 | 0.08 | 273 | 3.53 | <0.001 | *** |
| sT26: Invading:aT 32 | 0.68 | 0.53 | 0.83 | 0.07 | 273 | 9.18 | <0.001 | *** |
| sT22/32: Invading:aT 32 | 0.56 | 0.41 | 0.70 | 0.08 | 273 | 7.30 | <0.001 | *** |
| sT32: Invading:aT 32 | 0.32 | 0.16 | 0.48 | 0.08 | 273 | 3.96 | <0.001 | *** |

#### Supporting Table 9| Model selection table for mixed model on effect of salinity, temperature and biotic scenario on cell size.

The global model uses data for species' cell size measured in their selection temperature. Through model averaging to ensure that delta AICc was >2, the best model included selection salinity, species and biotic regime in full interaction. Selection temperature also affected cell size significantly, in interaction with species. Abbreviations are used as follows: Intc for intercept, sT for selection temperature (22ºC,26ºC, 32ºC or 22/32ºC), Inv, for biotic regime (colonising or invading), sp for species (*Ostreococcus* or *Chlamydomonas*), and Sal for the specie's selection salinity ('ancestral' for the selection salinity equal to its natural salinity, 'away' for evolution in a novel salinity). Further, df, for degrees of freedom, logLiK, for log likelihood, AICc as akaike information criterion for small sample sizes, delta as the difference in AICc between the best model and the other models, and weight, as the model weight (i.e. exp( -1/2 * delta) divided by the sum of this for all models) .+ denotes that a component was part of the model. We display the first ten out of 167 models.

| **Model** | **Intc** | **Sal** | **Inv** | **sT** | **sp** | **Sal Inv** | **Sal sT** | **Sal sp** | **Inv sT** | **Inv sp** | **sT sp** | **Sal Inv sT** | **Sal Inv sp** | **Sal sT sp** | **Inv sT sp** | **Sal Inv sT sp** | **df** | **logLik** | **AICc** | **delta** | **weight** |
|---|---|---|---|---|---|---|---|---|---|---|---|---|---|---|---|---|---|---|---|---|---|
| **1** | **9.58** | **+** | **+** | **+** | **+** | **+** | **NA** | **+** | **NA** | **+** | **+** | **NA** | **+** | **NA** | **NA** | **NA** | **16** | **-140.74** | **316.15** | **0.00** | **0.59** |
| 2 | 9.77 | + | + | + | + | + | + | + | NA | + | + | NA | + | + | NA | NA | 22 | -133.68 | 316.47 | 0.32 | 0.30 |
| 3 | 9.70 | + | + | + | + | + | + | + | NA | + | + | NA | + | NA | NA | NA | 19 | -137.67 | 317.13 | 0.98 | 0.04 |
| 4 | 9.79 | + | + | + | + | + | + | + | + | + | + | NA | + | + | NA | NA | 25 | -131.15 | 318.96 | 2.81 | 0.01 |
| 5 | 9.59 | + | + | + | + | + | NA | + | + | + | + | NA | + | NA | NA | NA | 19 | -138.94 | 319.67 | 3.52 | 0.05 |
| 6 | 9.72 | + | + | + | + | + | + | + | + | + | + | NA | + | NA | NA | NA | 22 | -135.64 | 320.39 | 4.23 | 0.00 |
| 7 | 9.84 | + | + | + | + | + | + | + | + | + | + | + | + | + | NA | NA | 28 | -128.98 | 322.42 | 6.27 | 0.00 |
| 8 | 9.68 | + | + | + | + | + | + | + | + | + | + | + | + | + | + | + | 34 | -120.89 | 322.57 | 6.42 | 0.00 |
| 9 | 9.77 | + | + | + | + | + | + | + | + | + | + | + | + | NA | NA | NA | 25 | -133.33 | 323.32 | 7.17 | 0.00 |
| 10 | 9.81 | + | + | + | + | + | + | + | + | + | + | NA | + | + | + | NA | 28 | -129.47 | 323.41 | 7.25 | 0.00 |

Supporting Table 10| Model averaging output on analysis of mixed effects model identified from Supporting Table 9 **(cell size)**

p values <0.05 are in bold. Abbreviations of model parameters are as in Supporting Table 9 above.

| | **Estimate** | **Std Error** | **Adj Std Error** | **z value** | **p value** |
|---|---|---|---|---|---|
| Sal Away, sp *Chlamydomonas,* sT 22ºC, inv colonising | 9.67364 | 0.1925 | 0.19371 | 49.938 | **2.00E-16** |
| Sal Ancestral | -0.0563 | 0.23004 | 0.23131 | 0.243 | 0.8077 |
| Inv Invading | -1.13043 | 0.16128 | 0.16562 | 6.825 | **2.00E-16** |
| sT 26ºC | -1.49682 | 0.17649 | 0.17781 | 8.418 | **2.00E-16** |
| sT 32ºC | -2.10741 | 0.20893 | 0.2101 | 10.031 | **2.00E-16** |
| sT 22/32ºC | -0.92306 | 0.30705 | 0.30797 | 2.997 | **0.00272** |
| sp *Ostreococcus* | -7.89203 | 0.21523 | 0.21679 | 36.404 | **2.00E-16** |
| Sal Ancestral: Inv Invading | 1.14811 | 0.20087 | 0.20242 | 5.672 | **2.00E-16** |
| Sal Ancestral: sp *Ostreococcus* | 0.00439 | 0.24826 | 0.25002 | 0.018 | 0.98599 |
| Inv Invading: sp *Ostreococcus* | 1.07776 | 0.20347 | 0.20504 | 5.256 | **1.00E-07** |
| sT 26ºC : sp *Ostreococcus* | 0.99962 | 0.22324 | 0.22484 | 4.446 | **8.80E-06** |
| sT 32ºC : sp *Ostreococcus* | 1.55636 | 0.24725 | 0.24881 | 6.255 | **2.00E-16** |
| sT 22/32ºC : sp *Ostreococcus* | 0.40055 | 0.30915 | 0.31045 | 1.29 | 0.19698 |
| Sal Home : Inv Invading: sp *Ostreococcus* | -1.24901 | 0.26399 | 0.26602 | 4.695 | **2.70E-06** |
| Sal Ancestral: sT 26ºC | 0.02936 | 0.24579 | 0.24755 | 0.119 | 0.90559 |
| Sal Ancestral: sT 32ºC | 0.3241 | 0.26026 | 0.26202 | 1.237 | 0.21611 |
| Sal Ancestral: sT 22/32ºC | 0.64961 | 0.31525 | 0.31683 | 2.05 | **0.04033** |
| Sal Ancestral: sT 26ºC: sp *Ostreococcus* | 0.2875 | 0.35618 | 0.35898 | 0.801 | 0.42321 |
| Sal Ancestral: sT 32ºC: sp *Ostreococcus* | -0.35577 | 0.36976 | 0.37267 | 0.955 | 0.33976 |
| Sal Ancestral: sT 22/32ºC: sp *Ostreococcus* | -0.62852 | 0.375 | 0.37795 | 1.663 | 0.09632 |

#### Supporting Table 11| Model selection for mixed effects model on effects of selection regimes on growth rates in decomposed samples

The global model considers data for samples in their selection temperature. The best model included species and the former role of the species in the mixed cultures in interaction, but not selection temperature. Abbreviations are used as follows: Intc for intercept, sT for selection temperature (22ºC,26ºC, 32ºC or 22/32ºC), sp for species (*Ostreococcus* or *Chlamydomonas*), and was for the specie's role in the mixed culture (resident or invader). Further, df, for degrees of freedom, logLiK, for log likelihood, AICc as akaike information criterion for small sample sizes, delta as the difference in AICc between the best model and the other models, and weight, as the model weight (i.e. exp( -1/2 * delta) divided by the sum of this for all models) .+ denotes that a component was part of the model. We display the first ten out of 20 possible models.

| **Model** | **Intc** | **Sp** | **sT** | **Was** | **sp * sT** | **sp * was** | **sT * was** | **spsT * was** | **df** | **logLik** | **AICc** | **delta** | **weight** |
|---|---|---|---|---|---|---|---|---|---|---|---|---|---|
| **1** | **1.12** | **+** | **NA** | **+** | **NA** | **+** | **NA** | **NA** | **7** | **78.99** | **-142.70** | **0.00** | **0.96** |
| 2 | 1.15 | + | + | + | NA | + | NA | NA | 10 | 82.17 | -139.76 | 2.94 | 0.02 |
| 3 | 1.17 | + | + | + | NA | + | + | NA | 13 | 83.68 | -136.92 | 5.78 | 0.01 |
| 4 | 1.15 | + | + | + | + | + | NA | NA | 13 | 82.20 | -133.96 | 8.74 | 0.00 |
| 5 | 1.17 | + | + | + | + | + | + | NA | 16 | 83.70 | -128.52 | 14.18 | 0.00 |
| 6 | 1.15 | + | + | + | + | + | + | + | 19 | 86.30 | -124.60 | 18.10 | 0.00 |
| 7 | 1.05 | + | NA | + | NA | NA | NA | NA | 6 | 62.20 | -111.46 | 31.24 | 0.00 |
| 8 | 1.08 | + | + | + | NA | NA | NA | NA | 9 | 64.43 | -108.76 | 33.94 | 0.00 |
| 9 | 1.10 | + | + | + | NA | NA | + | NA | 12 | 65.46 | -103.16 | 39.53 | 0.00 |
| 10 | 1.08 | + | + | + | + | NA | NA | NA | 12 | 64.44 | -101.13 | 41.57 | 0.00 |

Supporting Table 12| Model output on analysis of mixed effects model identified from Supporting Table 11 **(growth and decomposition)** 'Df' is degrees of freedom; abbreviations for factors in the model are as described above , t-values are the ratios of parameter estimates and standard errors. p values <0.05 are in bold.

| | Estimate | Std. Error | DF | t-value | p-value |
|---|---|---|---|---|---|
| sp (Chlamydomonas) was (resident) | 1.12 | 0.022 | 46 | 50.47 | **0.001** |
| was (invader) | -0.38 | 0.031 | 46 | -12.12 | **0.001** |
| sp (*Ostreococcus*) | -0.25 | 0.031 | 10 | -7.858 | **0.001** |
| was (invader) : sp (*Ostreococcus*) | 0.28 | 0.0441 | 46 | 6.21 | **0.001** |

#### Supporting Table 13| Model selection table for mixed model on effect of salinity, temperature and biotic scenario on net primary productivity in evolved and decomposed samples

The global model uses data for decomposed samples after 200 generations of evolution. The best model included selection temperature, species and whether the sample was a coloniser, a mixed, or decomposed (former resident or invader) sample, but not in interaction. Abbreviations are used as follows: Intc for intercept, sT for selection temperature (22ºC,26ºC, 32ºC or 22/32ºC), was for the role of the species during co-culture or the composition of the sample (invader , resident, mixed sample, coloniser) and sp for species (*Ostreococcus* or *Chlamydomonas*). Further, df, for degrees of freedom, logLiK, for log likelihood, AICc as akaike information criterion for small sample sizes, delta as the difference in AICc between the best model and the other models, and weight, as the model weight (i.e. exp( -1/2 * delta) divided by the sum of this for all models) .+ denotes that a component was part of the model. We display the first ten out of 20 models.

| **Model** | **Intc** | **Sp** | **sT** | **was** | **sp* sT** | **sp * was** | **sT * was** | **sp * St * was** | **df** | **logLik** | **AICc** | **delta** | **weight** |
|---|---|---|---|---|---|---|---|---|---|---|---|---|---|
| **1** | **0.16** | **+** | **+** | **+** | **NA** | **NA** | **NA** | **NA** | **10.00** | **85.39** | **-149.83** | **0.00** | **0.72** |
| 2 | 0.18 | NA | + | + | NA | NA | NA | NA | 9.00 | 82.50 | -146.22 | 3.61 | 0.12 |
| 3 | 0.15 | + | + | + | NA | NA | + | NA | 19.00 | 93.21 | -144.96 | 4.87 | 0.06 |
| 4 | 0.16 | + | + | + | + | NA | NA | NA | 13.00 | 85.92 | -144.22 | 5.61 | 0.04 |
| 5 | 0.15 | + | + | + | NA | + | NA | NA | 13.00 | 85.76 | -143.90 | 5.93 | 0.04 |
| 6 | 0.18 | NA | + | + | NA | NA | + | NA | 18.00 | 90.17 | -141.25 | 8.58 | 0.01 |
| 7 | 0.15 | + | + | + | + | NA | + | NA | 22.00 | 93.77 | -138.87 | 10.96 | 0.00 |
| 8 | 0.15 | + | + | + | NA | + | + | NA | 22.00 | 93.62 | -138.57 | 11.26 | 0.00 |
| 9 | 0.15 | + | + | + | + | + | NA | NA | 16.00 | 86.28 | -138.12 | 11.71 | 0.00 |
| 10 | 0.15 | + | + | + | + | + | + | NA | 25.00 | 94.18 | -132.28 | 17.55 | 0.00 |

Supporting Table 14| Model output on analysis of mixed effects model identified from Supporting Table 13 **(net primary production and decomposition).** 'Df' is degrees of freedom, t-values are the ratios of parameter estimates and standard errors. p values <0.05 are in bold. Abbreviations of model parameters are as in Supporting Table 13 above.

| | Estimate | Std.Error | DF | t-value | p-value |
|---|---|---|---|---|---|
| was (coloniser) sp Chlamydomonas sT 22ºC | 0.156 | 0.028 | 173 | 5.486 | **<0.001** |
| was (invader) | -0.107 | 0.031 | 173 | -3.492 | **0.001** |
| was (full sample) | 0.070 | 0.031 | 173 | 2.260 | **0.025** |
| was (resident) | 0.053 | 0.031 | 173 | 1.720 | 0.087 |
| sp *Ostreococcus* | 0.053 | 0.022 | 173 | 2.376 | **0.019** |
| sT 22/32ºC | 0.222 | 0.031 | 59 | 7.192 | **<0.001** |
| sT 26ºC | 0.214 | 0.031 | 173 | 6.793 | **<0.001** |
| sT 32ºC | -0.069 | 0.031 | 173 | -2.205 | **0.029** |

Supporting Table 15| t-test results for comparing growth rate μ in samples in mono-culture after decomposition and during co-culture (short term). There was no significant difference (across temperature regimes) between growth after decomposition when samples had only grown in co-culture for 14 days and growth in mono-culture.

| **Welch Two Sample t-test** | |
|---|---|
| t = 0.6489, df = 189.3, p-value = 0.5172 | |
| alternative hypothesis: true difference in means is not equal to 0 | |
| 95 percent confidence interval: | |
| -0.036 | 0.072 |
| mean growth in mono-culture short | |
| 0.87 | |
| mean growth after decomposition short | |
| 0.853 | |

#### Supporting Table 16| Model selection table for mixed model on effect of salinity, temperature and biotic scenario on biomass

The global model uses data for species' biomass measured at selection temperature. The best model included selection temperature, species and biotic regime in full interaction. Abbreviations are used as follows: Intc for intercept, sT for selection temperature (22ºC,26ºC, 32ºC or 22/32ºC), InvTr, for invasion treatment including the resident species (colonising, invading, and being invaded), sp for species (*Ostreococcus* or *Chlamydomonas*). Further, df, for degrees of freedom, logLiK, for log likelihood, AICc as akaike information criterion for small sample sizes, delta as the difference in AICc between the best model and the other models, and weight, as the model weight (i.e. exp( -1/2 * delta) divided by the sum of this for all models) .+ denotes that a component was part of the model. We display the first ten out of 167 models.

| Model | Intc | sp | sT | InvTr | sp sT | sp InvTr | stT InvTr | sp InvTr sT | df | logLik | AICc | delta | weight |
|---|---|---|---|---|---|---|---|---|---|---|---|---|---|
| **1** | **21.07** | + | + | + | + | + | + | + | **35** | **-180.76** | **445.14** | **0.00** | **1.00** |
| 2 | 21.42 | + | + | + | + | + | + | NA | 26 | -295.82 | 650.87 | 205.73 | 0.00 |
| 3 | 21.08 | + | + | + | + | NA | + | NA | 23 | -304.21 | 660.02 | 214.88 | 0.00 |
| 4 | 21.06 | + | + | + | + | + | NA | NA | 17 | -315.63 | 668.28 | 223.15 | 0.00 |
| 5 | 20.77 | + | + | + | + | NA | NA | NA | 14 | -323.74 | 677.53 | 232.39 | 0.00 |
| 6 | 21.43 | + | + | + | NA | + | + | NA | 23 | -324.05 | 699.70 | 254.56 | 0.00 |
| 7 | 21.15 | + | + | + | NA | + | NA | NA | 14 | -335.59 | 701.22 | 256.08 | 0.00 |
| 8 | 21.24 | + | + | + | NA | NA | + | NA | 20 | -329.83 | 703.86 | 258.73 | 0.00 |
| 9 | 20.96 | + | + | + | NA | NA | NA | NA | 11 | -341.36 | 705.99 | 260.85 | 0.00 |
| 10 | 19.67 | + | NA | + | NA | + | NA | NA | 11 | -388.62 | 800.50 | 355.37 | 0.00 |

Supporting Table 17| Model output on analysis of mixed effects model identified from Supporting Table 16 **(biomass).** 'Df' is degrees of freedom; t-values are the ratios of parameter estimates and standard errors. p values <0.05 are in bold. Abbreviations of model parameters are as in Supporting Table 16 above.

| | Value | Std.Error | DF | t-value | p-value |
|---|---|---|---|---|---|
| InvTr Ancestral salinity (colonising) , sp *Chlamydomonas*, sT 22ºC | 21.07 | 0.22 | 110 | 94.045 | **0.0002** |
| InvTr (novel salinity, colonising) | -4.23 | 0.43 | 45 | -9.862 | **0.0002** |
| InvTr (novel salinity, invading) | -4.50 | 0.34 | 34 | -13.102 | **0.0002** |
| InvTr (resident) | 1.32 | 0.32 | 34 | 4.170 | **0.0002** |
| sT 26ºC | -0.42 | 0.32 | 110 | -1.320 | 0.1896 |
| sT 32ºC | -4.01 | 0.32 | 110 | -12.651 | **0.0002** |
| sT 22/32ºC | -1.19 | 0.32 | 110 | -3.749 | **0.0002** |
| sp *Ostreococcus* | -3.31 | 0.32 | 34 | -10.449 | **0.0002** |
| InvTr (novel salinity, colonising) : sT 26ºC | 0.61 | 0.58 | 110 | 1.051 | 0.2957 |
| InvTr (novel salinity, invading): sT 26ºC | 1.11 | 0.47 | 34 | 2.383 | 0.0229 |
| InvTr (resident) : sT 26ºC | -1.90 | 0.45 | 34 | -4.235 | **0.0002** |
| InvTr (novel salinity, colonising) : sT 32ºC | 3.12 | 0.64 | 110 | 4.841 | **0.0002** |
| InvTr (novel salinity, invading): sT 32ºC | 2.53 | 0.47 | 34 | 5.418 | **0.0002** |
| InvTr (resident) : sT 32ºC | -0.20 | 0.46 | 34 | -0.445 | 0.6589 |
| InvTr (novel salinity, colonising) : sT 22/32ºC | 1.61 | 0.61 | 110 | 2.648 | **0.0093** |
| InvTr (novel salinity, invading): sT 22/32ºC | -1.60 | 0.50 | 34 | -3.214 | **0.0029** |
| InvTr (resident) : sT 22/32ºC | -1.06 | 0.45 | 34 | -2.373 | **0.0234** |
| InvTr (novel salinity, colonising) :sp *Ostreococus* | 1.49 | 0.53 | 45 | 2.785 | **0.0078** |
| InvTr (novel salinity, invading) :sp *Ostreococus* | 2.96 | 0.47 | 34 | 6.283 | **0.0002** |
| InvTr (resident) :sp *Ostreococus* | -3.03 | 0.45 | 34 | -6.765 | **0.0002** |
| sT 26ºC: sp *Ostreococcus* | -3.51 | 0.45 | 34 | -7.840 | **0.0002** |
| sT 32ºC: sp *Ostreococcus* | 2.51 | 0.46 | 34 | 5.496 | **0.0002** |
| sT 22/32ºC: sp *Ostreococcus* | -1.60 | 0.45 | 34 | -3.578 | **0.0011** |
| InvTr (novel salinity, colonising) : sT 26ºC: sp *Ostreococcus* | 1.65 | 0.73 | 110 | 2.258 | **0.0259** |
| InvTr (novel salinity, invading) : sT 26ºC: sp *Ostreococcus* | -1.63 | 0.65 | 34 | -2.512 | **0.0169** |
| InvTr (resident) : sT 26ºC: sp *Ostreococcus* | 6.84 | 0.63 | 34 | 10.800 | **0.0002** |
| InvTr (novel salinity, colonising) : sT 32ºC: sp *Ostreococcus* | -5.04 | 0.82 | 110 | -6.137 | **0.0002** |
| InvTr (novel salinity, invading) : sT 32ºC: sp *Ostreococcus* | -6.82 | 0.66 | 34 | -10.389 | **0.0002** |
| InvTr (resident) : sT 32ºC: sp *Ostreococcus* | 0.52 | 0.65 | 34 | 0.807 | 0.425 |
| InvTr (novel salinity, colonising) : sT 22/32ºC: sp *Ostreococcus* | 0.22 | 0.75 | 110 | 0.291 | 0.7714 |
| InvTr (novel salinity, invading) : sT 22/32ºC: sp *Ostreococcus* | 2.89 | 0.68 | 34 | 4.275 | **0.0001** |
| InvTr (resident) : sT 22/32ºC: sp *Ostreococcus* | 3.50 | 0.63 | 34 | 5.527 | **0.0002** |
